# Noise-induced hearing loss enhances Ca^2+^-dependent spontaneous bursting activity in lateral cochlear efferents

**DOI:** 10.1101/2025.01.07.631771

**Authors:** Hui Hong, Laurence O Trussell

## Abstract

Exposure to loud and/or prolonged noise damages cochlear hair cells and triggers downstream changes in synaptic and electrical activity in multiple brain regions, resulting in hearing loss and altered speech comprehension. It remains unclear however whether or not noise exposure also compromises the cochlear efferent system, a feedback pathway in the brain that fine-tunes hearing sensitivity in the cochlea. We examined the effects of noise-induced hearing loss on the spontaneous action potential (AP) firing pattern in mouse lateral olivocochlear (LOC) neurons. This spontaneous firing exhibits a characteristic burst pattern dependent on Ca^2+^ channels, and we therefore also examined the effects of noise-induced hearing loss on the function of these and other ion channels. The burst pattern was sustained by an interaction between inactivating Ca^2+^ currents contributed largely by L-type channels, and steady outward currents mediated by Ba^2+^-sensitive inwardly-rectifying and two-pore domain K^+^ channels. One week following exposure to loud broadband noise, hearing thresholds were significantly elevated, and the duration of AP bursts was increased, likely as a result of an enhanced Ca^2+^ current. Additional effects of noise-induced hearing loss included alteration of Ca^2+^-dependent inactivation of Ca^2+^ currents and a small elevation of outward K^+^ currents. We propose that noise-induced hearing loss enhances efferent activity and may thus amplify the release of neurotransmitters and neuromodulators (i.e., neuropeptides), potentially altering sensory coding within the damaged cochlea.

**Significance Statement:** Although the effects of noise-induced hearing loss on the auditory afferent system have been extensively studied, little is known about its impact on the auditory efferent system, which modulates hearing sensitivity via feedback from the brain. Additionally, while Ca^2+^ channels are related to numerous neurological diseases, their involvement in auditory disorders is underexplored. This study bridges these gaps by examining Ca^2+^ channel-driven spontaneous burst firing in lateral olivocochlear (LOC) neurons, the most numerous auditory efferent neurons. Noise-induced hearing loss differentially affects Ca^2+^ channel subtypes by increasing high-voltage activated currents that further prolong burst firing and suggesting altered intracellular Ca^2+^ signaling. These significant changes in LOC firing behavior may profoundly impact their downstream targets in the cochlea.

## Introduction

Noise-induced hearing loss (NIHL) is a major health concern in the world and is closely associated with other auditory disorders such as tinnitus and hyperacusis (1–3). In the periphery, excessive noise exposure leads to hair cell death and auditory nerve degeneration (4, 5). Such attenuation of sound input to the brain has prolonged effects on the central auditory system. In the cochlear nucleus – the first relay station in the auditory brainstem, a variety of neurons increase their spontaneous firing rate and excitability, apparently to compensate for diminished input activity (6, 7). For example, fusiform cells in the dorsal cochlear nucleus relay their hyperactivity to the inferior colliculus in the midbrain and may thus mediate tinnitus (8–10). Similar compensatory strategy has also been proposed for the auditory cortex, leading to enhancement in the strength of its descending projections (11). Moreover, all these changes in electrical and synaptic properties of auditory neurons following noise exposure are closely related to alterations in gene expression (12, 13).

Despite decades of research on the impact of NIHL on the brain, a significant gap in knowledge exists in the auditory efferent system, a feedback system from the brain to the ear that is believed to protect and modify hearing sensitivity in response to diverse acoustic environments (14). Lateral olivocochlear (LOC) neurons, the most numerous auditory efferent neurons, are located in the lateral superior olive (LSO) of the brainstem (15, 16). They send axons primarily to the ipsilateral cochlea, where they synapse onto dendrites of spiral ganglion neurons that form the auditory nerve. By expressing a wide range of neurotransmitters and neuropeptides, LOC neurons may modulate the activity of spiral ganglion neurons, thereby influencing hearing sensitivity (14, 17, 18). Interestingly, recent studies found that noise-induced hearing loss upregulates dopamine and neuropeptide expression in LOC neurons, which could further inhibit the spiral ganglion neurons (19, 20). However, it is not clear how the electrical properties of LOC are affected in this prevalent disorder.

We recently discovered a unique infra-slow (∼0.1 Hz) spontaneous burst firing pattern in LOC neurons of juvenile and young adult mice *in vitro*, driven by a Ca^2+^-dependent intrinsic neuronal oscillator (21). Here, we report that such spontaneous activity and the underlying ion channel properties are significantly impacted by NIHL. High-voltage activated Ca^2+^ (largely L-type) current was increased one week after noise exposure, while low-voltage activated (T-type) Ca^2+^ current was unchanged. Ca^2+^ channels exhibited Ca^2+^-dependent inactivation (CDI), and this form of inactivation was also modified following noise exposure. Larger Ca^2+^ current led to a pronounced increase in burst duration in LOC neurons. In addition to burst initiation, we also examined the ion channel mechanisms for burst termination. Ba^2+-^sensitive two-pore domain K^+^ (K2P) channels and inwardly-rectifying K^+^ (K_ir_) channels likely function to facilitate the termination of bursts as Ca^2+^ current inactivates. We propose that efferent regulation is enhanced in the noise-exposed cochlea.

## Results

### Ca^2+^ channels are essential for burst initiation in LOC

All experiments were conducted on offspring from crosses between ChAT-IRES-Cre and Ai9 mice in order to visualize LOC neurons with tdTomato for brain slice recordings (see Materials and Methods) (21). Our previous study characterized the spontaneous burst firing activity in LOC neurons from juvenile and young adult mice using four techniques: cell-attached, whole-cell current-clamp, perforated patch recordings, and calcium imaging. All four methods yielded a similar burst frequency of approximately 0.1 Hz (21). Therefore, in this study, we chose to use cell-attached recordings, which allow us to monitor the activity of individual LOC neurons *in vitro* for over an hour. Consistent with our previous finding, Ca^2+^ channels play an essential role in burst initiation, as broad-spectrum Ca^2+^ channel blockers Cd^2+^ and Ni^2+^ (100 µM each) abolished the bursts in all of recorded neurons (burst frequency: control = 0.139 ± 0.028 Hz, Cd^2+^ and Ni^2+^ = 0 ± 0 Hz; n = 7 cells, W = -28, p = 0.016, Wilcoxon matched-pairs signed rank test), and washout of these blockers restored the patterned activity in 2 out of 7 neurons (**Figure 1**).

**Figure 1.**
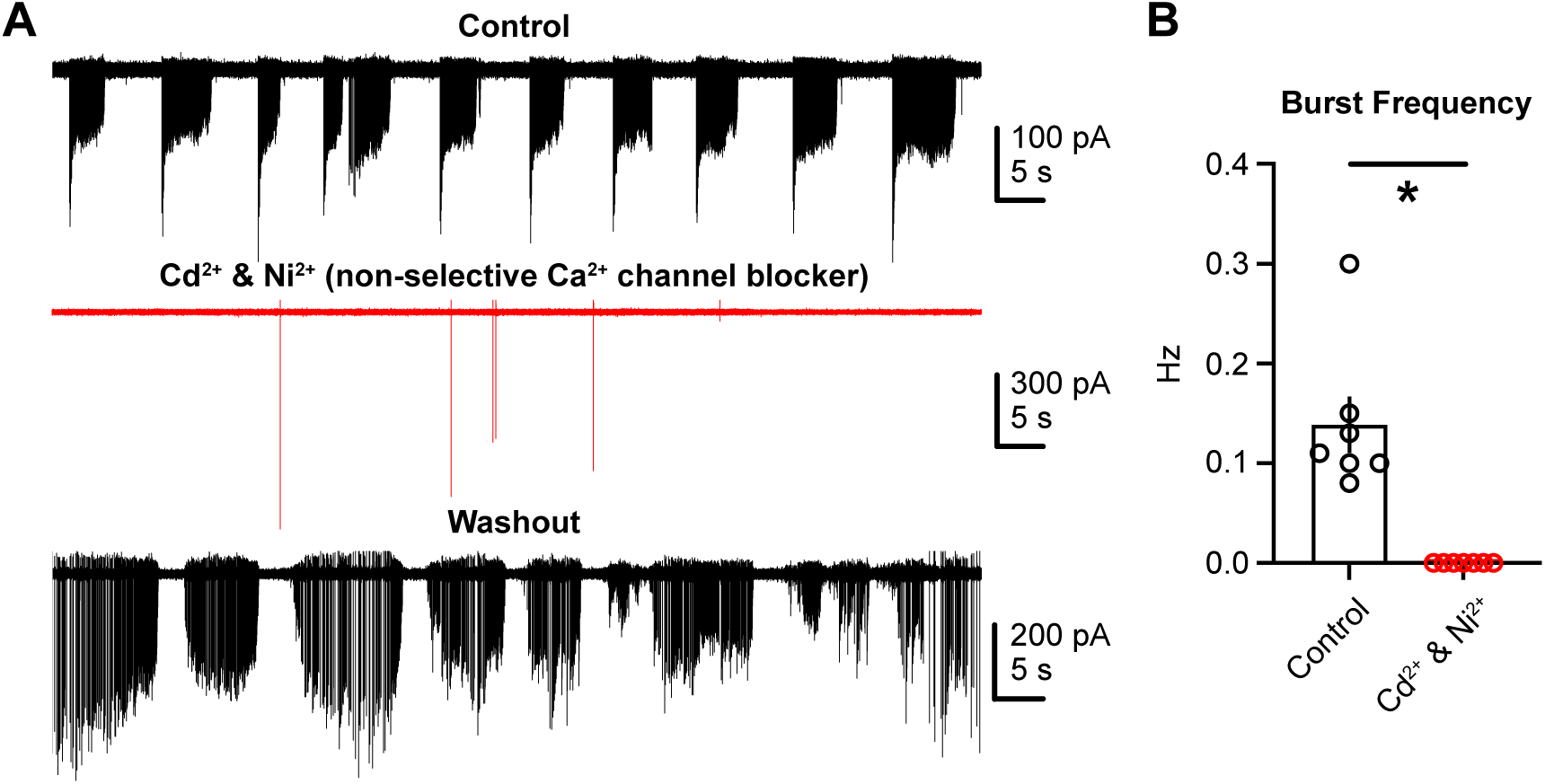
Ca^2+^ channels are essential for burst initiation in LOC neurons. (A) Representative 100-s cell-attached recordings in control, with bath application of Cd^2+^ and Ni^2+^ (100 µM each) and after washout. (B) Population data showing burst frequency in control and with Cd^2+^ and Ni^2+^. N = 7 cells. *p = 0.016. Error bars, SEM.

### K^+^ channels that are necessary for burst termination in LOC

We next examined the ion channels required for burst termination. As there is Ca^2+^ influx associated with the bursts (21), we anticipated that Ca^2+^-activated K^+^ channels provided repolarizing drive essential for burst termination. Previous snRNA-seq study has also demonstrated the mRNA expression of these channels (**Figure S1A**) (19). However, spontaneous burst firing persisted with bath application of iberiotoxin (100 nM), a selective antagonist of large-conductance Ca^2+^-activated K^+^ channels (BK), without significant changes in burst frequency (**Figure 2A1-A2**; burst frequency: control = 0.074 ± 0.016 Hz, iberiotoxin = 0.074 ± 0.012 Hz; n = 5 cells, t(4) < 0.0001, p > 0.9999, paired t-test). Similar results were obtained when apamin (200 nM) was used to block small-conductance Ca^2+^-activated K^+^ channels (SK, **Figure 2B1-B2**; burst frequency: control = 0.100 ± 0.009 Hz, apamin = 0.103 ± 0.018 Hz; n = 4 cells, t(3) = 0.103, p = 0.925, paired t-test). BK and SK channels did exhibit roles in regulating the inter-spike interval (ISI) of action potentials (APs) during the bursts. **Figure 2A3 and 2B3** show the distribution of ISI from all of recorded neurons under different conditions. With iberiotoxin, the peak of ISI distribution shifts significantly from 38 to 28 ms, corresponding to an increase in spiking frequency from 26 to 36 Hz (**Figure 2A3**; Kolmogorov-Smirnov D = 0.142, p < 0.0001, Kolmogorov-Smirnov test). These results indicate that BK channels reduce the spiking frequency during the bursts, likely due to their contribution to spike afterhyperpolarization (22–24). In contrast, SK channels facilitated spiking: with apamin, although the peak of ISI distribution remained at ∼32 ms, the distribution was skewed towards longer ISIs (**Figure 2B3**, arrowhead; Kolmogorov-Smirnov D = 0.221, p < 0.0001, Kolmogorov-Smirnov test).

**Figure 2.**
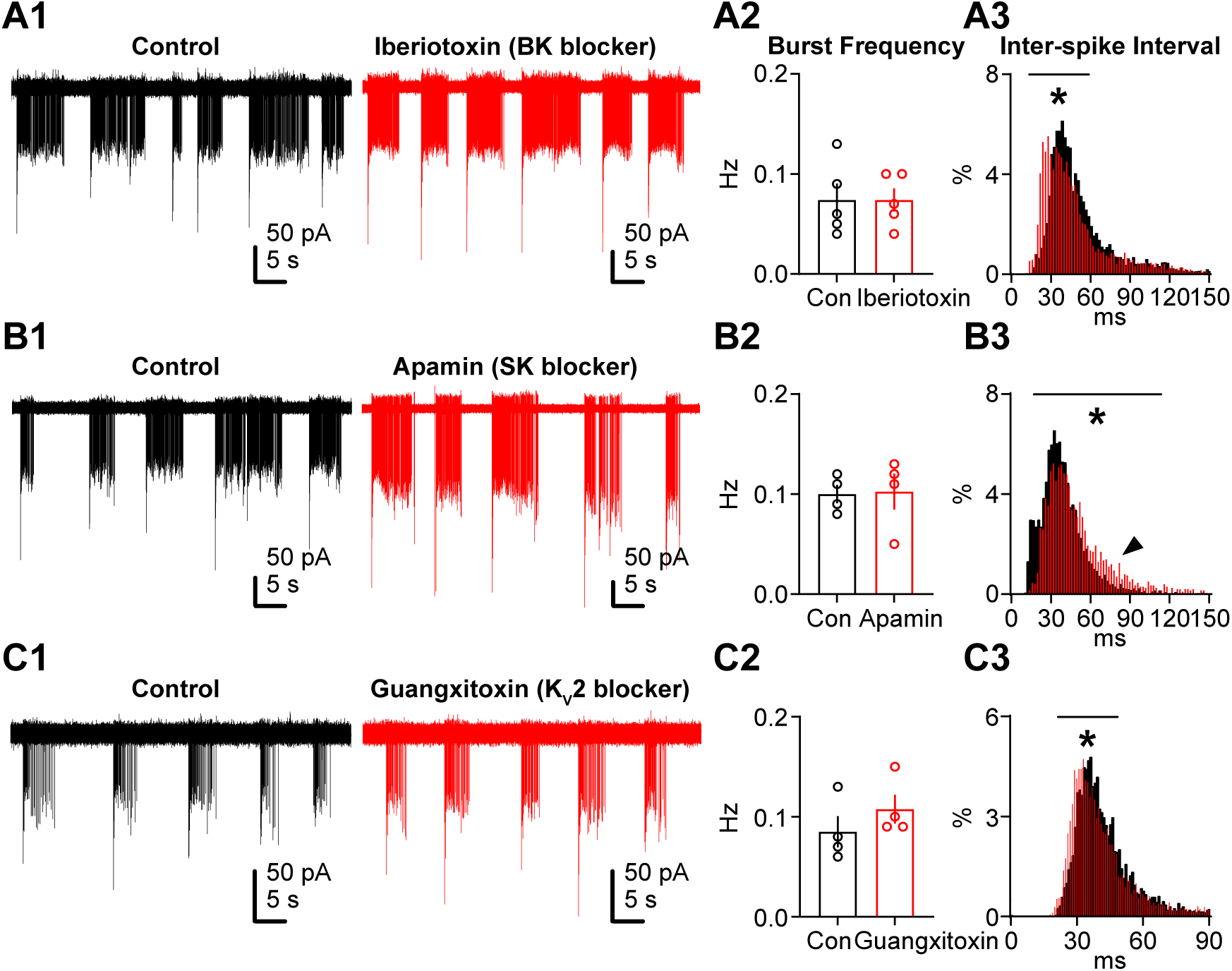
Ca^2+^-activated BK, SK and K_V_2 channels do not contribute to burst termination in LOC neurons. (A1-C1) Representative 60-s cell-attached recordings in control, with bath application of iberiotoxin (A1, 100 nM), apamin (B1, 200 nM) or guangxitoxin (C1, 100 nM). Experiments using iberiotoxin and guangxitoxin had 0.5 mg/mL bovine serum albumin (BSA) in the bath. (A2-C2) Population data showing burst frequency in control and with bath application of three different drugs. (A3-C3) Overlaid histograms showing frequency distribution of inter-spike intervals (ISIs) from all neurons in control and drug-treated conditions. Arrowhead in B3 points to the skewed tail towards longer ISIs with apamin. Iberiotoxin: N = 5 cells, *p < 0.0001; apamin: N = 4 cells, *p < 0.0001; guangxitoxin: N = 4 cells, *p < 0.0001 (Kolmogorov-Smirnov tests). Error bars, SEM.

We next examined whether voltage-gated K^+^ channels played a role in burst termination. As *Kcnb2* mRNA encoding the Kv2.2 channel is expressed in LOC neurons (**Figure S1B**) (19), we tested the effect of guangxitoxin, a selective antagonist for K_V_2 channels. However, guangxitoxin (100 nM) had no significant effect on burst frequency (**Figure 2C1-C2**; burst frequency: control = 0.085 ± 0.016 Hz, guangxitoxin = 0.108 ± 0.014 Hz; n = 4 cells, W = 10, p = 0.125, Wilcoxon matched-pairs signed rank test). Rather, blocking K_V_2 channels led to a slight but significant left shift in ISI distribution (**Figure 2C3**; Kolmogorov-Smirnov D = 0.090, p < 0.0001, Kolmogorov-Smirnov test), again indicating a subtle role of K_V_2 channels in regulating spiking frequency.

*Kcnq1*, *Kcnq3* and *Kcnq5* mRNA encoding K_V_7.1, K_V_7.3 and K_V_7.5 were also detected in LOC neurons (**Figure S1C**) (19), and heterogeneous effects were observed upon application of the K_V_7 channel antagonist, XE-991 (10 µM). A minority of neurons (3 out of 13) became silent with XE-991 (**Figure 3A1**), while the majority maintained their burst firing pattern similar to the control (**Figure 3A2-A3**; burst frequency: control = 0.132 ± 0.016 Hz, XE-991 = 0.119 ± 0.024 Hz; n = 13 cells, t(12) = 0.552, p > 0.9999, Bonferroni post hoc test), indicating that K_V_7 channels do not terminate bursts, and suggesting that there may be heterogeneity in ion channel distribution among the population of LOC neurons. We then applied a non-selective K^+^ channel antagonist, tetraethylammonium (TEA, 10 mM). Spontaneous activity still persisted with TEA, but with obvious alterations in spiking pattern. Bursts of activity became shorter and contained a single onset spike followed by a plateau current, resembling depolarization block (**Figure 3A2**, inset, arrowheads). Additionally, burst frequency increased significantly with XE-991 and TEA (**Figure 3A3**; burst frequency: XE-991 & TEA = 0.299 ± 0.053 Hz; n = 9 cells, t(8) = 3.596, p = 0.021, Bonferroni post hoc test). Thus, TEA-sensitive channels are required to maintain steady spiking during a burst, but blocking these channels did not fully eliminate oscillatory activity. Finally, spontaneous activity was abolished by subsequent application of Ba^2+^ (0.1-2 mM; **Figure 3A2-A3**; burst frequency: XE-991, TEA & Ba^2+^ = 0 Hz; n = 7 cells, t(6) = 6.075, p = 0.003, Bonferroni post hoc test). These results indicate that the K^+^ channels essential for burst termination in LOC neurons are sensitive to Ba^2+^, pointing to two candidates: K2P and K_ir_ channels (25).

**Figure 3.**
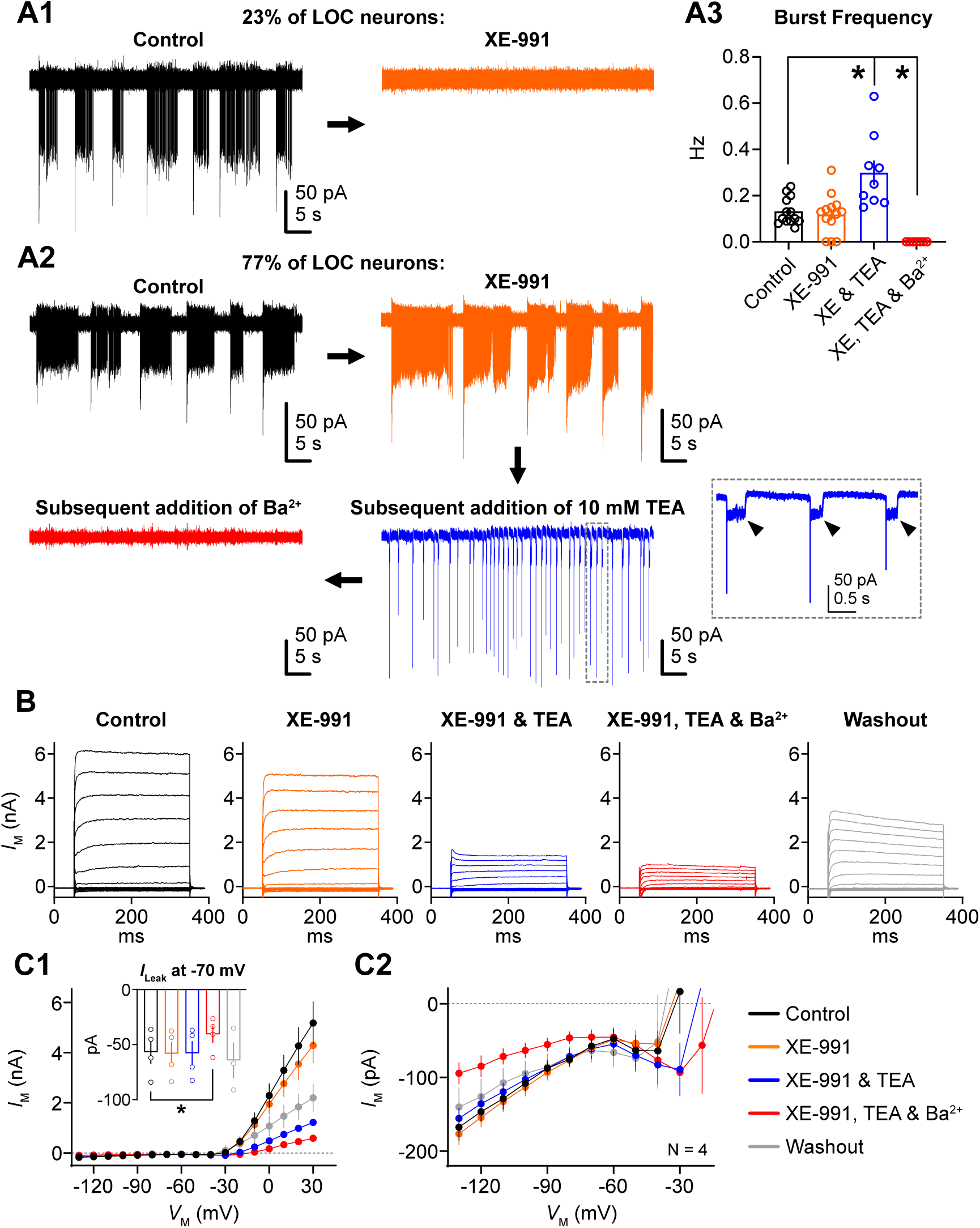
TEA-insensitive but Ba^2+^-sensitive K^+^ channels are essential for burst termination in LOC neurons. (A1-A2) Representative 60-s cell-attached recordings showing heterogenous responses to bath application of XE-991 (10 µM): while the minority of neurons (23%, A1) became silent with XE-991, the majority (77%, A2) remained burst firing. In A2, TEA (10 mM) and Ba^2+^ (0.1-2 mM) were subsequently applied in the bath. Arrows indicate the order of drug application. The area highlighted by dashed lines in the blue trace is enlarged in the inset. Arrowheads point to the steady-state current immediately after the single onset spikes. (A3) Population data showing burst frequency in control and drug-treated conditions. N = 13, 13, 9 and 7 cells from left to right. *p = 0.021 (Control vs XE & TEA) and 0.003 (control vs XE, TEA & Ba^2+^). (B) Representative current traces from an LOC neuron in response to voltage steps from -130 to +30 mV in 10-mV steps, under control conditions, following the application of various drugs, and after washout. Holding voltage is -70 mV. (C1-C2) Population data showing the I-V relation of the steady-state current under different conditions. C2 enlarges the small current between -130 and -20 mV. Inset shows the population data of leak current (*I*_Leak_) at -70 mV. Leak subtraction was not applied. N = 4 cells. Error bars, SEM.

Whole-cell voltage-clamp experiments were conducted to examine the amount of K2P and K_ir_ current in LOC neurons. Individual neurons were held at -70 mV, and then a series of voltage steps from -130 to +30 mV were applied in 10-mV steps. **Figure 3B** shows current responses from a representative LOC neuron exposed to a sequence of drug applications. Current-voltage relationship under different conditions is shown in **Figure 3C1**, and the small current changes between -130 and -20 mV are expanded in **Figure 3C2**. XE-991 slightly reduced the outward current, while 10 mM TEA had the most pronounced effect, reducing the outward current by ∼72% at +30 mV from the XE-991-only condition to the XE-991 plus TEA condition. However, such a large reduction in outward K^+^ current was apparently not sufficient to halt spontaneous bursts (see **Figure 3A2**). Further reduction in current (at +30 mV, by 51%) was observed with the application of 1 mM Ba^2+^ (**Figure 3C1**, red trace). This current likely represents a combination of K2P and K_ir_ currents. However, analysis of the snRNA-seq database revealed the expression of *Kcnj3*, which encodes K_ir_3.1, a G-protein gated “strong” inward rectifier with minimal outward current at depolarized voltages (**Figure S1D**) (19, 26). Thus, the majority of this Ba^2+^-sensitive outward current is likely attributable to voltage-independent K2P currents. Washout of all the drugs partially recovered the outward current (**Figure 3B-C**, grey traces). In contrast to the outward current, inward current at the membrane voltages more negative than -90 mV (the reversal potential of K^+^ channels) was only sensitive to Ba^2+^ (**Figure 3C2**, red trace), consistent with the pharmacological properties of K2P and K_ir_ current (25). The Ba^2+^-induced block was reversible upon washout (**Figure 3C2**, grey trace).

We noticed that Ba^2+^ had effects beyond blocking K^+^ currents. During voltage-clamp experiments, LOC neurons were held at -70 mV, with a slightly negative holding current (*I*_Leak_, control = -56.86 ± 10.53 pA). The absolute amplitude of this current was not affected by XE-991 or TEA but was significantly reduced by subsequent application of Ba^2+^, suggesting the presence of a Ba^2+^-sensitive tonic cationic current in LOC neurons near rest. (**Figure 3C1**, inset; XE-991, TEA & Ba^2+^ = -40.66 ± 7.62 pA; n = 4 cells, t(3) = 3.52, p = 0.04, paired t-test).

In contrast to the clear expression of K_ir_3.1 channels, the mRNA expression of K2P channels appears less definitive. For example, **Figure S1E** shows apparent expression of *Kcnk2*, which encodes TREK-1 channels, in neonatal LOC neurons but ambiguous expression in mature neurons (19). To clarify this, immunohistochemical experiments were conducted to examine the expression of TREK-1 channel proteins in P25 LOC neurons. We used an anti-KCNK2 (TREK-1) antibody that has been validated on TREK-1 knockout mice (27). In the cerebellum, TREK-1 channels are extensively expressed in the molecular layer, in stark contrast to their minimal presence in the granule cell layer (**Figure 4A**). This protein expression pattern is consistent with a previous report on *Kcnk2* mRNA in cerebellum (28). In LOC neurons, TREK-1 staining is much less prominent than in cerebellum, but nevertheless appears on the plasma membrane as bright puncta (**Figure 4B-D**, arrowheads). TREK-1 expression is also observed in the cytoplasm as numerous speckles, but it is absent in the cell nucleus (**Figure 4B-D**). In a separate experiment, immunostaining was performed with secondary antibody only (i.e., Alexa 488). We observed minimal green background and no bright puncta in LOC neurons (**Figure 4E**).

**Figure 4.**
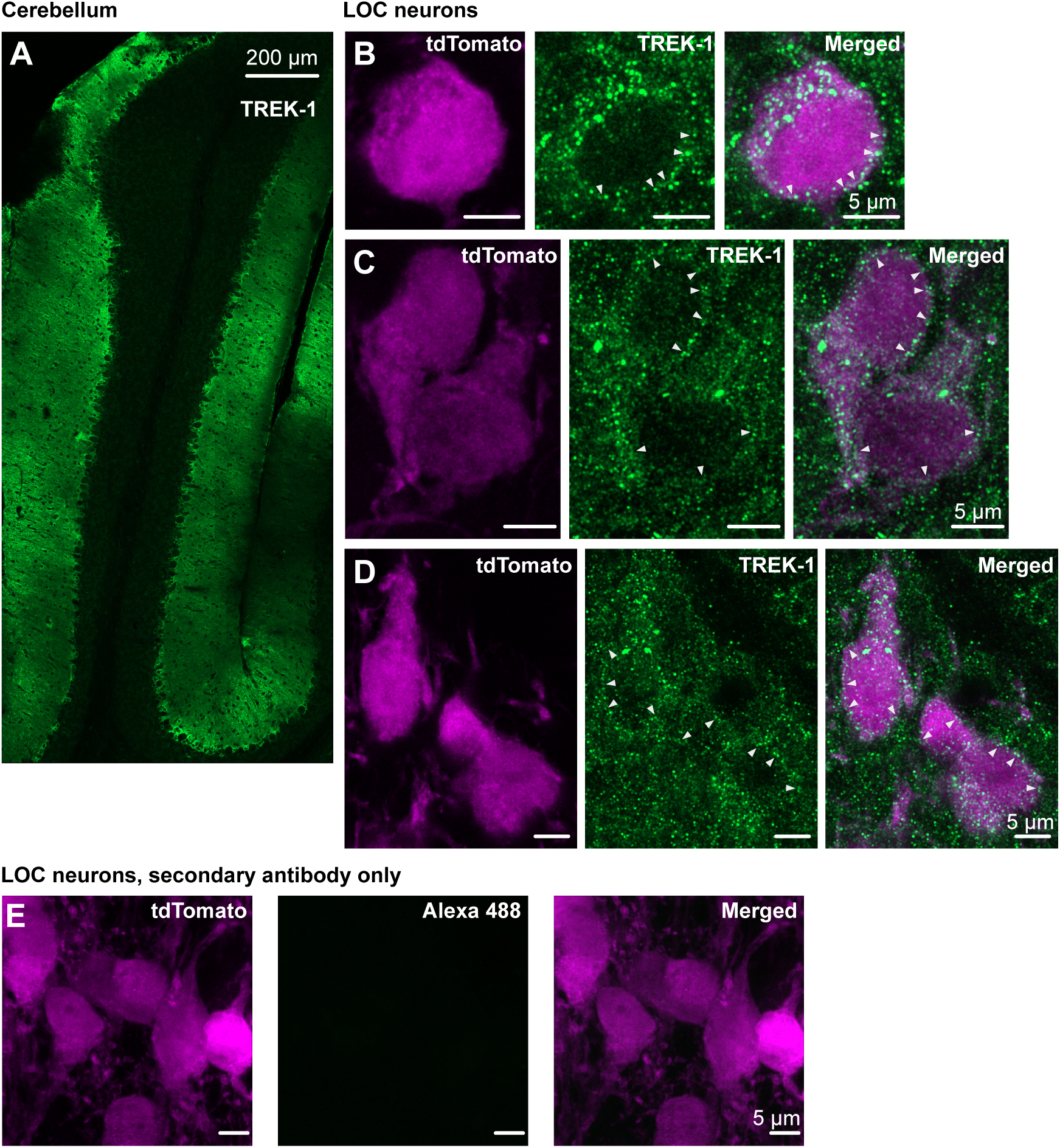
TREK-1 channels are expressed in LOC neurons. (A) Low-magnification confocal images showing extensive expression of TREK-1 in the molecular layer but not in the granule cell layer of the cerebellum. (B-D) High-magnification images displaying the expression of TREK-1 in tdTomato-positive LOC neurons, with TREK-1 staining appearing as puncta and speckles in both the plasma membrane (arrowheads) and cytoplasm, but not in the cell nucleus. (E) High-magnification images from immunohistochemical experiments where only the secondary antibodies (i.e., Alexa 488) were applied, omitting the primary anti-KCNK2 (TREK-1) antibodies. Minimal green background is present. Images in these experiments were captured using the same laser settings on the LSM980 and adjusted to similar levels of contrast and brightness in Fiji as in B-D.

### Ca^2+^ channel properties in LOC neurons

K2P channels carry tonic K^+^ current (29). We tested whether a tonic, hyperpolarizing current, combined with a voltage-sensitive Ca^2+^ current, would alone be sufficient to create voltage oscillations in LOC neurons that resemble the physiological measurements. Whole-cell current-clamp recordings were made using a CsCl-based internal solution to block the majority of K^+^ channels, while extracellular TTX (0.5 µM) blocked Na^+^ channels. When a small negative bias current (average -9±3.2 pA, range 0 to -30 pA) was injected through the recording electrode, LOC neurons exhibited oscillations at a frequency remarkably similar to that observed with the K-gluconate-based internal solution (**Figure 5A-B**; K^+^ internal = 0.103 ± 0.012 Hz, n = 15 cells, Cs^+^ internal = 0.097 ± 0.014 Hz, n = 10 cells; t(23) = 0.302, p = 0.765, Student t-test).

**Figure 5.**
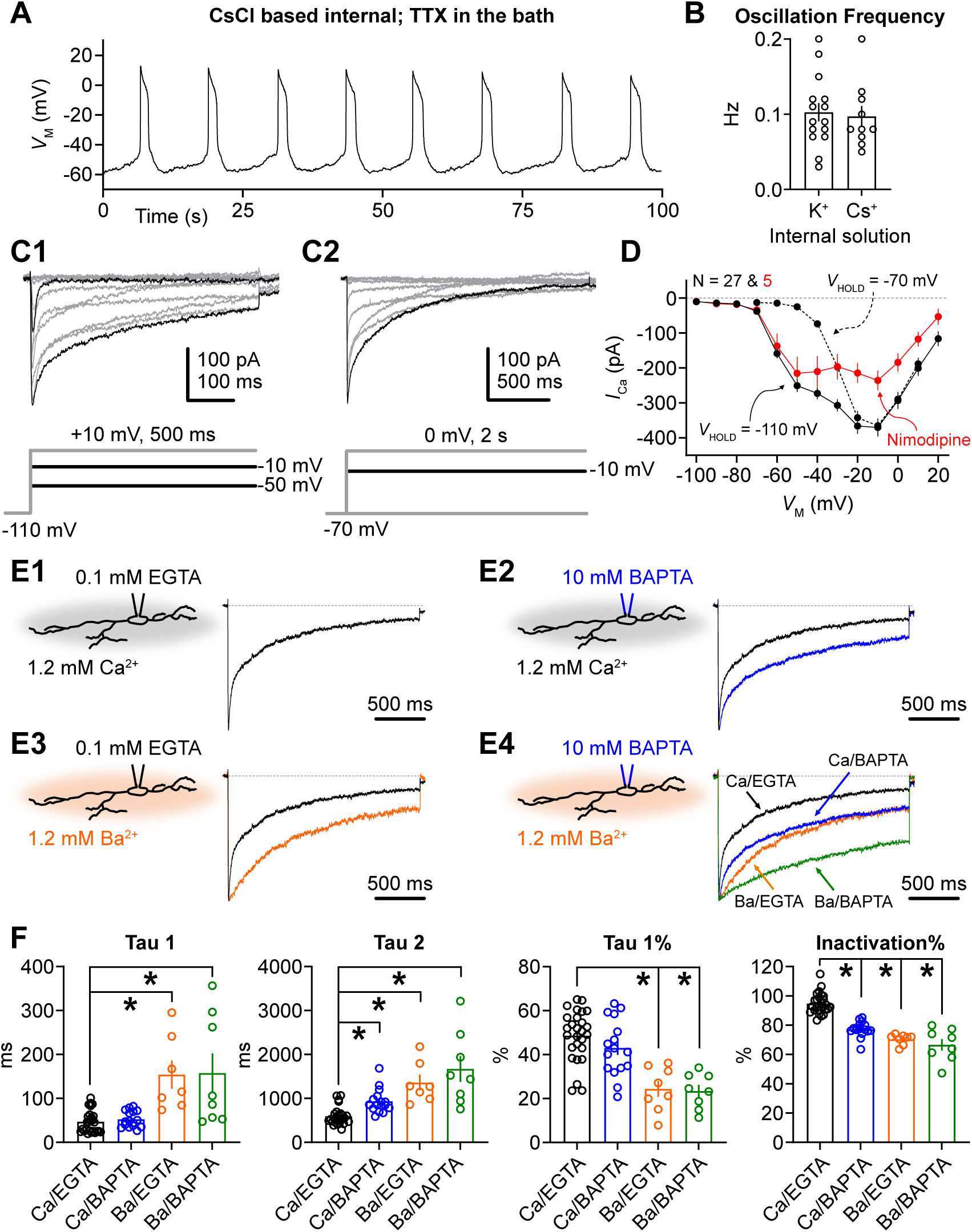
Ca^2+^-dependent Ca^2+^ channel inactivation (CDI) in LOC neurons. (A) Representative spontaneous depolarizations recorded from an LOC neuron using a CsCl-based internal pipette solution to block the majority of K^+^ channels, along with TTX (0.5 µM) in the bath to block Na^+^ channels. NBQX (10 µM), MK-801 (10 µM), strychnine (0.5 µM) and gabazine (10 µM) were also included to block synaptic activity. No bias current was applied to this neuron. (B) Population data showing the oscillation frequency of LOC neurons recorded with regular K^+^-based internal solution or specific CsCl-based internal solution. When using K^+^-based solution, no drugs were applied in the bath. N = 15 cells (K^+^ internal) and 10 cells (Cs^+^ internal). (C1-C2) Representative Ca^2+^ current traces from an LOC neuron in response to two different voltage-clamp protocols shown below the trace: C1 with a holding voltage (*V*_HOLD_) of -110 mV and 500-ms voltage steps, C2 with a *V*_HOLD_ of -70 mV and 2-s voltage steps. P/8 leak subtraction was applied to the neuron. TEA (100 mM), 4-AP (3-4 mM), TTX (0.5 µM), NBQX (10 µM), MK-801 (10 µM), strychnine (0.5 µM) and gabazine (10 µM) were applied in the bath to isolate Ca^2+^ current. (D) Population data showing the I-V relation of the peak Ca^2+^ current with different *V*_HOLD_. Nimodipine (10 µM) was applied in the bath to examine the amount of L-type Ca^2+^ current. N = 27 cells (control) and 5 cells (nimodipine). (E1-E4) Representative normalized Ca^2+^ current evoked from -70 to -20 mV under the combination of following conditions: EGTA (0.1 mM) or BAPTA (10 mM) in the internal solution, Ca^2+^ (1.2 mM) or Ba^2+^ (1.2 mM) in the external solution. Schematics depicting these conditions are shown to the left of each trace. (F) Population data showing the parameters of inactivation kinetics (tau 1, tau 2 and tau 1%) and percent inactivation of Ca^2+^ current from the peak to the end of 2-s recording window (Inactivation%) under four conditions. Left to right in each figure, greater CDI attenuation; N = 26, 16, 7 and 8 cells for tau 1, tau 2 and tau 1%; N = 27, 16, 8 and 8 cells for inactivation%. Tau 1: *p = 0.015 (Ca/EGTA vs Ba/EGTA) and 0.003 (Ca/EGTA vs Ba/BAPTA). Tau 2: *p < 0.0001 (Ca/EGTA vs Ca/BAPTA), *p = 0.005 (Ca/EGTA vs Ba/EGTA), *p < 0.0001 (Ca/EGTA vs Ba/BAPTA). Tau 1%: *p = 0.0006 (Ca/EGTA vs Ba/EGTA), *p < 0.0001 (Ca/EGTA vs Ba/BAPTA). Inactivation%: *p < 0.0001 (Ca/EGTA vs Ca/BAPTA), *p = 0.008 (Ca/EGTA vs Ba/EGTA), *p < 0.0001 (Ca/EGTA vs Ba/BAPTA). Error bars, SEM.

The persistence of voltage oscillations under conditions in which Ca^2+^ channels provided the majority of voltage-sensitive conductance prompted us to examine Ca^2+^ channel activation and inactivation. Two voltage-clamp protocols were applied to evoke Ca^2+^ currents. Individual neurons were held at -110 mV and given a series of voltage steps from -100 to +20 mV in a step of 10 mV for 500 ms (**Figure 5C1**). Next, the holding voltage was changed to -70 mV and voltage steps from -70 to +10 mV were applied to the neuron for 2 s (**Figure 5C2**). P/8 leak subtraction was applied during these protocols. K^+^ and Na^+^ channel blockers, along with synaptic receptor blockers were included in the bath to isolate Ca^2+^ currents (see Materials and Methods). The current-voltage relationship for Ca^2+^ current is shown in **Figure 5D**. With holding voltage of -110 mV, two distinct phases of activation were observed: a low-voltage activated Ca^2+^ current that was absent when holding voltage was changed to -70 mV; a high-voltage activated Ca^2+^ current that was partially sensitive to nimodipine (10 µM), an L-type Ca^2+^ channel antagonist. The low-voltage activated Ca^2+^ current was not blocked by nimodipine (**Figure 5D**), and its voltage dependence and rapid inactivation suggests that it is mediated by T-type Ca^2+^ channels. Therefore, LOC neurons have T- and L-type Ca^2+^ channels.

Given that T-type channels inactivate fully at the depolarized potentials associated with seconds-long burst activity, we focused on the L-type current. A typical feature of L-type Ca^2+^ channel is Ca^2+^-dependent Ca^2+^ channel inactivation (CDI) (30). Reasoning that such inactivation may contribute to the termination of the spike bursts and slow voltage oscillations, we next explored CDI by applying a single voltage step from -70 to -20 mV for 2 s (**Figure 5E1**). To attenuate CDI, we used an internal solution containing a rapid Ca^2+^ chelator, BAPTA (10 mM), and recorded the Ca^2+^ current at -20 mV. EGTA (0.1 mM) was used as control. A double exponential fit was applied to the inactivation phase of Ca^2+^ current, allowing us to calculate the fast and slow decay time constants (tau 1 and tau 2, respectively), along with their percent contribution to the total decay (see Materials and Methods). In **Figure 5E2**, Ca^2+^ currents recorded with EGTA or BAPTA were normalized and overlaid to illustrate the differences in the decay phase. A significant increase in tau 2 was observed when switching the intracellular buffer from EGTA to BAPTA (**Figure 5F**; Ca/EGTA = 594.8 ± 39.8 ms, n = 26 cells, Ca/BAPTA = 935.9 ± 68.0 ms, n = 16 cells; U = 56, p < 0.0001, Mann-Whitney U). As a result, the percent inactivation in Ca^2+^ current amplitude from the peak to the end of 2-s recording window decreased from ∼95% with EGTA to ∼77% with BAPTA (**Figure 5F**; U = 2, p < 0.0001, Mann-Whitney U).

CDI can also be attenuated by substituting extracellular Ca^2+^ with equimolar Ba^2+^. In the presence of Ba^2+^, peak currents were 51% smaller than in equimolar (i.e., 1.2 mM) Ca^2+^-containing bath solutions (n = 8 cells) (31). When currents in Ba^2+^ were normalized to those in Ca^2+^ the differences in inactivation kinetics were readily apparent (**Figure 5E3**). Ba^2+^ not only increased tau 2 (**Figure 5F**; Ba/EGTA = 1,360 ± 176 ms, n = 7 cells; t(6) = 4.334, p = 0.005, paired t-test), but also prolonged tau 1 (Ca/EGTA = 32.28 ± 9.19 ms, Ba/EGTA = 154.10 ± 32.06 ms; n = 7 cells, t(6) = 3.355, p = 0.015, paired t-test) and reduced its percent contribution to the fit (i.e., tau 1%; Ca/EGTA = 47.51 ± 4.32%, Ba/EGTA = 24.33 ± 3.57%; t(8) = 5.916, p = 0.0006, paired t-test). In one neuron, the fraction of tau 1 was less than 10%, so a single-exponential fit was used to calculate the decay time constant for this neuron. Thus, tau 2 became more dominant in the decay phase when CDI was attenuated. The slower decay kinetics of Ba^2+^ current resulted in lower percent inactivation, at ∼70% (W = -36, p = 0.008, Wilcoxon matched-pairs signed rank test). Further attenuation of CDI was achieved by combining BAPTA and Ba^2+^ (**Figure 5E4 & 5F**). In summary, CDI in LOC neurons accounts for at least 30% of Ca^2+^ channel inactivation.

### Noise-induced hearing loss prolongs burst firing and alters Ca^2+^ current

Recent studies show that NIHL enhances dopamine and neuropeptide expression in LOC neurons (19, 20). The release pattern of neurotransmitters is closely related to the firing pattern of the neuron (32–34). Hence, we investigated whether NIHL changes spontaneous burst firing and the underlying ion channel properties in LOC neurons. Auditory brainstem responses (ABRs) were recorded in P20-22 mice to ensure normal hearing. One day after ABR testing, the mice were exposed to broadband noise (4-50 kHz) at 110 dB SPL for 2 hours. 6-7 days after noise exposure, ABR experiments were conducted again on the same mice to assess the degree of hearing loss. Brain slices were then collected from deafened mice within a week after the second ABR testing (see Materials and Methods). **Figure 6A** shows the representative ABR traces before and 1-week after noise exposure in response to 16 kHz pure tone. ABR signal was minimal after noise exposure even at the maximal sound intensity tested (90 dB SPL). ABR thresholds were elevated by 20-50 dB SPL from 4 to 32 kHz sound stimulation, indicating profound hearing loss (**Figure 6B**; at 16 kHz: pre-exposure = 27.08 ± 1.28 dB SPL, 1-week post-exposure = 78.67 ± 1.60 dB SPL, n = 60 ears; t(59) = 25.16, p < 0.0001, Bonferroni post hoc test). 2 weeks after noise exposure, ABR thresholds for low-frequency sound were recovered, while remained elevated for high-frequency sound, reflecting a permanent threshold shift (at 16 kHz: 2-week post-exposure = 75.00 ± 1.64 dB SPL, n = 8 ears; t(7) = 21.19, p < 0.0001, Bonferroni post hoc test compared to pre-exposure data). Cell-attached and whole-cell current-clamp recordings were conducted on noise-exposed LOC neurons and their age-matched controls to assess spontaneous burst firing pattern (**Figure 6C1-C2**). NIHL significantly increased the burst duration while simultaneously reducing the burst frequency (**Figure 6D-E**; burst duration: control = 3.80 ± 0.33 s, n = 37 cells, noise-exposed = 8.50 ± 0.94 s, n = 29 cells; U = 191, p < 0.0001; burst frequency: control = 0.113 ± 0.009 Hz, noise-exposed = 0.064 ± 0.004 Hz; U = 188.5, p < 0.0001, Mann-Whitney U for both comparisons).

**Figure 6.**
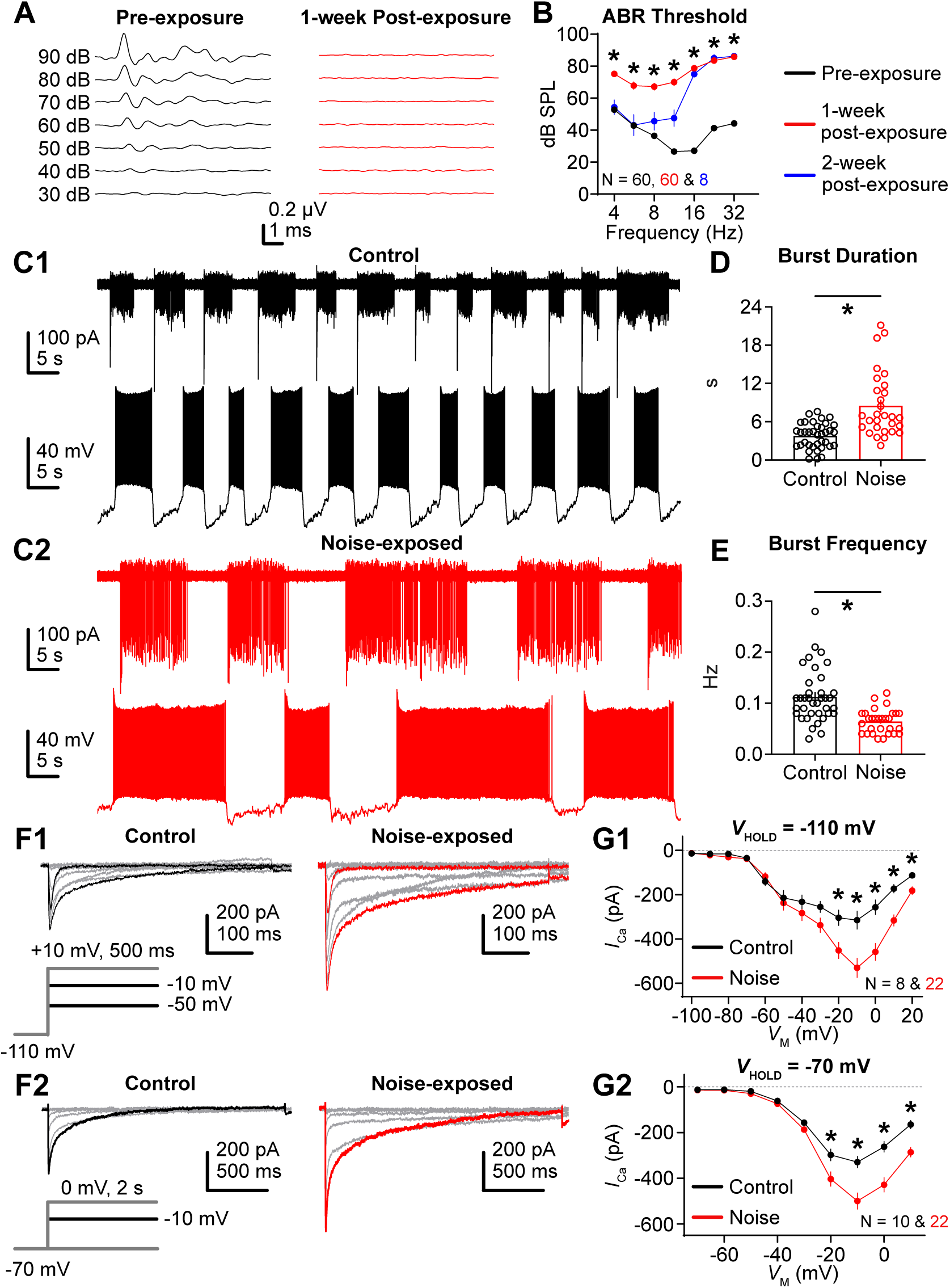
Noise-induced hearing loss alters electrical properties of LOC neurons. (A) Representative auditory brainstem responses (ABRs) to a 16-kHz pure tone from a ChAT-Cre/tdTomato mouse recorded before and one week after exposure to broadband noise (4-50 kHz) at 110 dB SPL for 2 hours. The numeric values to the left of the ABR traces indicate the level of the pure tone. (B) Population data showing the ABR thresholds at 4, 5.6, 8, 11.3, 16, 22.6 and 32 kHz from pre-, 1-week and 2-week post-exposure groups. *p < 0.0001 for all frequencies between pre- and 1-week post-exposure groups. N = 60 ears for pre- and 1-week post-exposure groups, 8 ears for 2-week post-exposure group. (C1-C2) Representative 100-s cell attached (upper panel) and whole-cell current-clamp (lower panel) recordings from age-matched control and noise-exposed LOC neurons. (D-E) Population data showing burst duration and frequency in control and noise-exposed groups. Note that cell-attached and current-clamp data were pooled. N = 37 cells (control) and 29 cells (noise-exposed). *p < 0.0001 for both D and E. (F1-F2) Representative Ca^2+^ current traces from control and noise-exposed neurons in response to two different voltage-clamp protocols shown below the trace: F1 with a holding voltage (*V*_HOLD_) of -110 mV and 500-ms voltage steps, F2 with a *V*_HOLD_ of -70 mV and 2-s voltage steps. P/8 leak subtraction was applied to the neurons. (G1-G2) Population data showing the I-V relation of the peak Ca^2+^ current with different *V*_HOLD_. G1: N = 8 cells (control) and 22 cells (noise-exposed); *p at the peak = 0.01. G2: N = 10 cells (control) and 22 cells (noise-exposed); *p at the peak = 0.002. Error bars, SEM.

The high-voltage activated Ca^2+^ current was also altered by NIHL. **Figure 6F1-F2** show the representative Ca^2+^ current traces from control and noise-exposed neurons with holding voltage at -110 mV (**Figure 6F1**) or -70 mV (**Figure 6F2**). **Figure 6G1-G2** show the corresponding current-voltage relationship. An average of 56% increase in the absolute amplitude of Ca^2+^ current was observed at membrane voltages of -20 mV and above (at -10 mV: control = -329.2 ± 25.1 pA, n = 10 cells, noise-exposed = -500.0 ± 35.6 pA, n = 22 cells; U = 36, p = 0.0018, Mann-Whitney U). This indicates that NIHL enhances the high-voltage activated Ca^2+^ current without affecting low-voltage activated Ca^2+^ current. Furthermore, we observed no changes in the passive membrane properties (i.e., input resistance, membrane capacitance and tau) between control and noise-exposed groups (**Figure S2A**). A slight but significant increase in K^+^ current was observed between -20 and +10 mV following noise exposure (**Figure S2B-D**; at +10 mV: control = 3.21 ± 0.24 nA, n = 20 cells, noise-exposed = 3.96 ± 0.26 nA, n = 21 cells; U = 129, p = 0.035, Mann-Whitney U). Interestingly, we also observed an increase in inward current at -50 and -40 mV, which could reflect the aforementioned larger Ca^2+^ current, or could reveal additional changes in Na^+^ current, as no Ca^2+^ channel or Na^+^ channel antagonists were used in this experiment (**Figure S2E**; at = -40 mV: control = -52.84 ± 5.73 pA, noise-exposed = -75.39 ± 7.23 pA; t(39) = 2.43, p = 0.02, Student t-test).

Ca^2+^ channel inactivation was explored following noise exposure, specifically examining the role of CDI as assessed by comparing cells recorded with BAPTA (10 mM) vs EGTA (0.1 mM). **Figure 7A1-A2** show the representative Ca^2+^ current traces from normal-hearing control and noise-exposed groups at -20 mV. Tau 1 and tau 2 were calculated under different conditions (**Figure 7B**). Surprisingly, the expected elongation of tau 2 with BAPTA as in control neurons was no longer observed in noise-exposed neurons (**Figure 7B**, right; noise/EGTA = 664.3 ± 40.2 ms, n = 20 cells, noise/BAPTA = 592.2 ± 71.1 ms, n = 9 cells; t(67) = 0.833, p > 0.9999, Bonferroni post hoc test). As a result, tau 2 with BAPTA in noise-exposed neurons is significantly smaller than that in controls (control/BAPTA = 935.9 ± 68.0 ms, n = 16 cells; t(67) = 3.826, p = 0.002, Bonferroni post hoc test).

**Figure 7.**
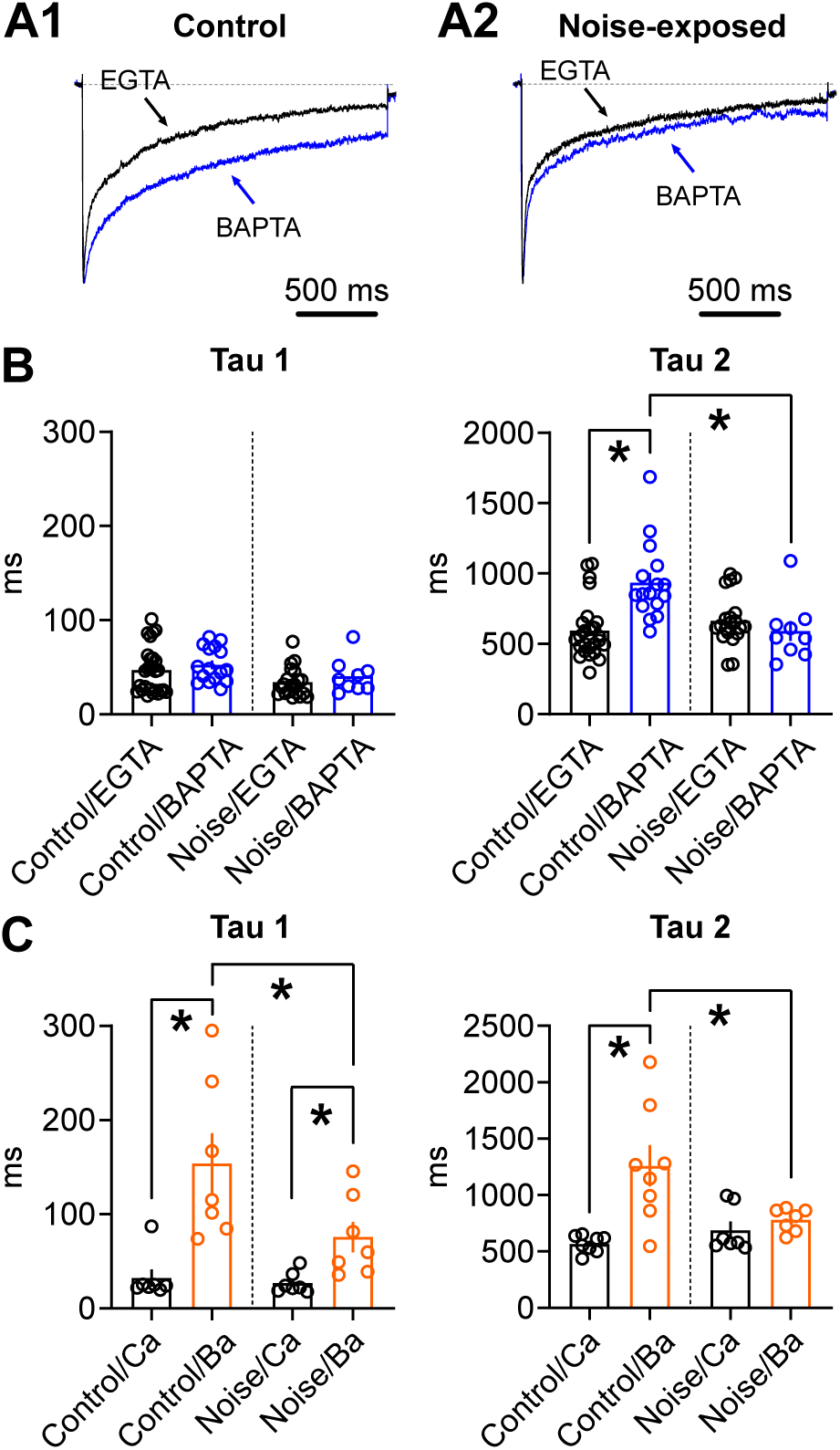
CDI properties in LOC neurons are compromised by noise-induced hearing loss. (A1-A2) Representative normalized Ca^2+^ current evoked from -70 to -20 mV in control and noise-exposed LOC neurons, with EGTA (0.1 mM) or BAPTA (10 mM) in the internal pipette solution. (B-C) Population data showing changes in tau 1 and tau 2 of inactivation under different conditions. B: N = 26, 16, 20 and 9 cells from left to right in each panel; tau 2: *p < 0.0001 (control/EGTA vs control/BAPTA), *p = 0.002 (control/BAPTA vs noise/BAPTA). C: N = 7, 7, 7 and 7 cells from left to right for tau 1; N = 8, 8, 7 and 7 cells from left to right for tau 2; tau 1: *p = 0.015 (control/Ca vs control/Ba), 0.027 (noise/Ca vs noise/Ba), 0.049 (control/Ba vs noise/Ba); tau 2: *p = 0.01 (control Ca vs control/Ba), 0.04 (control/Ba vs noise/Ba). Error bars, SEM.

Similar results were observed when Ba^2+^ was used to attenuate CDI in noise-exposed LOC neurons. Tau 2 did not change when switching from Ca^2+^ to Ba^2+^ (**Figure 7C**, right; noise/Ca = 688.2 ± 76.4 ms, noise/Ba = 779.8 ± 38.7 ms; n = 7 cells, t(6) = 1.495, p = 0.186, paired t-test), and thus it is significantly smaller than tau 2 with Ba^2+^ in control group (control/Ba = 1259.0 ± 183.4 ms, n = 8 cells; U = 10, p = 0.04, Mann-Whitney U). NIHL had an additional effect on tau 1. Although Ba^2+^ still lengthened tau 1 in noise-exposed neurons, the increase was less pronounced – 178% compared to a 381% increase in control neurons (**Figure 7C**, left; noise/Ca = 26.96 ± 4.25 ms, noise/Ba = 75.81 ± 16.10; t(6) = 2.914, p = 0.027, paired t-test). This resulted in significantly smaller tau 1 with Ba^2+^ in noise-exposed neurons than in controls (control/Ba = 154.10 ± 32.06 ms; n = 7 cells, t(12) = 2.182, p = 0.049, Student t test). Overall, our data indicate that with NIHL, the contribution of CDI to the process of inactivation was reduced. This was especially true for the slow phase of inactivation (i.e., tau 2). However, it is noteworthy that even with minimal CDI, noise-exposed neurons still exhibited profound Ca^2+^ current inactivation (**Figure 7B**). These data suggest that enhancement in voltage-dependent inactivation of Ca^2+^ channels may compensate for the loss of CDI (see Discussion).

### Computational modeling

Although Ca^2+^ channels are required for spontaneous burst firing in LOC neurons, and NIHL leads specifically to increases in high-voltage activated current, it is not clear whether this increase in Ca^2+^ current is responsible for prolonged bursts. To test this possibility, we built a minimal, Hodgkin-Huxley-type model of LOC neurons. This single-compartment model contained multiple ionic currents to capture the features of spontaneous spike burst activity. Model parameters are shown in Table 1 (see Materials and Methods). **Figure 8A1** shows the control simulation of spontaneous burst firing in model LOC neuron. The conductance of high-voltage activated Ca^2+^ channel (gCa = 0.8 mS/cm^2^) was derived from voltage-clamp recordings shown in **Figure 5D**. The modeled neuron had an average burst duration of 5.35 s, and a burst frequency of 0.12 Hz, resembling our experimental data (**Figure 6C1, 6D & 6E**).

**Figure 8.**
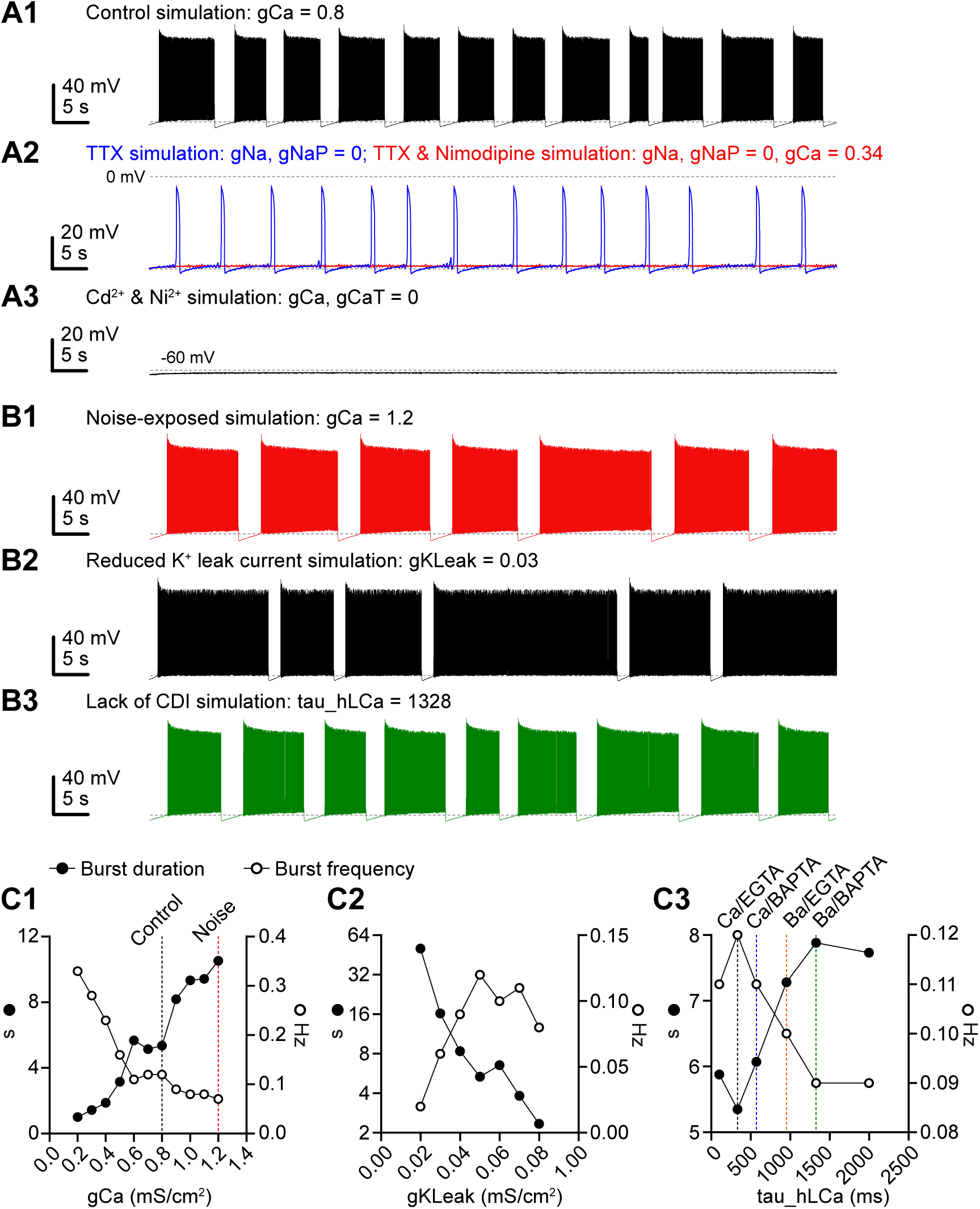
Model LOC neuron exhibits prolonged bursts with increased Ca^2+^ current. (A1) Control simulation of spontaneous burst firing with high-voltage activated Ca^2+^ conductance (gCa) set at 0.8 mS/cm^2^, based on experimental data from Figure 5D. Dashed line, -60 mV. (A2) To simulate the application of TTX, transient and persistent Na^+^ conductances were set to 0. Simulation of subsequent nimodipine application was achieved by reducing gCa to 0.34 mS/cm^2^, as per Figure 5D. Upper dashed line, 0 mV; lower dashed line, -60 mV. (A3) To simulate the application of Cd^2+^ and Ni^2+^, as in Figure 1, both gCa and low-voltage activated Ca^2+^ conductance (gCaT) were set to 0. Dashed line, -60 mV. (B1) To simulate noise-exposed condition, gCa was raised to 1.2 mS/cm^2^, according to Figure 6G2. (B2) To examine the role of K2P current in burst termination, its conductance (gKLeak) was reduced from 0.05 in control to 0.03 mS/cm^2^. (B3) To examine the role of CDI in burst termination, the inactivation time constant of high-voltage activated Ca^2+^ conductance (tau_hLCa) was raised from 333 to 1328 ms, corresponding to weighted tau shift from Ca/EGTA to Ba/BAPTA in Figure 5F. Dashed lines in B1-B3, -60 mV. (C1-C3) Changes in burst duration (left axis, filled circle) and burst frequency (right axis, open circle) with varying gCa, gKLeak and tau_hLCa. In C1, the black and red dashed lines represent the gCa in control and noise-exposed LOC neurons, respectively. In C3, the colored dashed lines indicate tau_hLCa under various CDI-attenuated conditions.

**Table 1.**
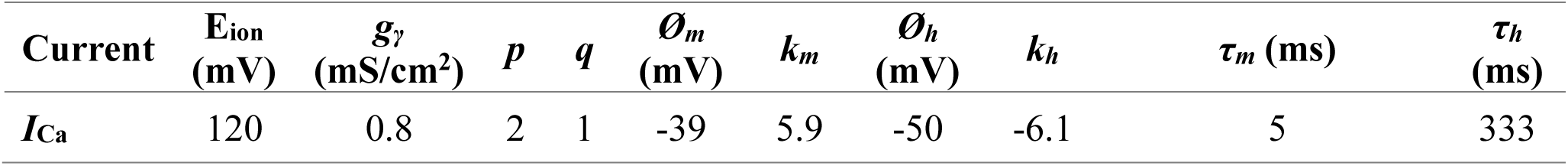
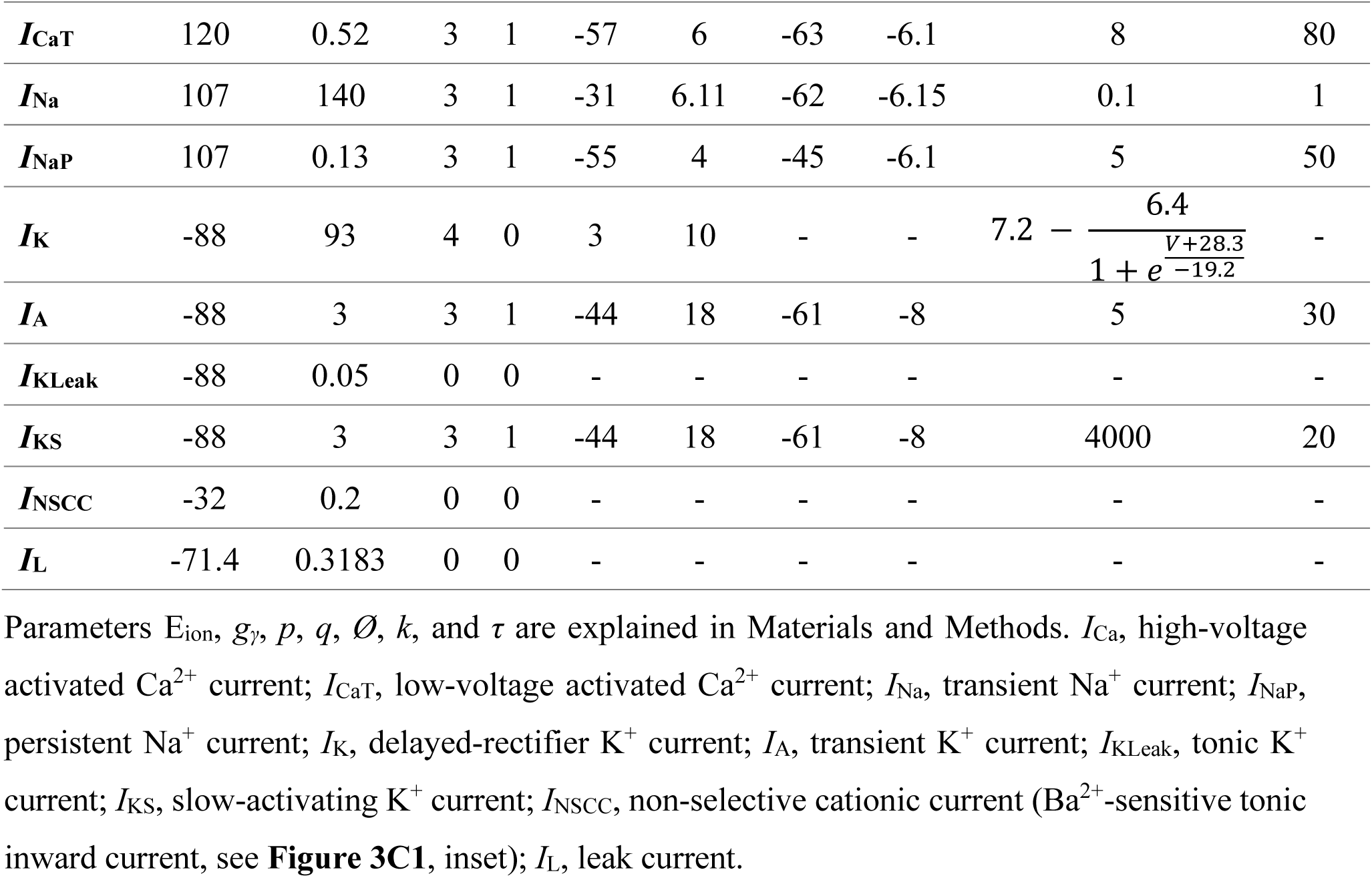
Ionic currents in the model LOC neuron.

In order to further test the model, we first set transient and persistent Na^+^ current (gNa and gNaP, respectively) at 0, which simulates the bath application of TTX. The model generated voltage oscillations without APs, and these were subsequently abolished by reducing high-voltage activated Ca^2+^ conductance from 0.8 to 0.34 mS/cm^2^, simulating bath application of nimodipine (**Figure 8A2**, also see **Figure 5D**). This result is consistent with our pharmacological experiments reported in Hong et al., 2022 (21). To simulate application of Cd^2+^ and Ni^2+^, we changed both high- and low-voltage activated Ca^2+^ conductance (gCa and gCaT) to 0. By doing so, model neuron failed to generate any patterned activity (**Figure 8A3**), consistent with cell-attached recordings shown in **Figure 1A**.

To assess the role of high-voltage activated Ca^2+^ current in regulating burst duration, we varied gCa and calculated burst duration and frequency in the model neuron. **Figure 8B1** shows spontaneous burst firing pattern when gCa = 1.2 mS/cm^2^, a value derived from voltage-clamp recordings in **Figure 6G2**, simulating increased Ca^2+^ current in noise-exposed LOC neurons. The average burst duration is 10.52 s and burst frequency is 0.07 Hz, closely matching the experimental data shown in **Figure 6C2, 6D and 6E**. Burst duration and frequency were then plotted as a function of varying gCa (**Figure 8C1**). We observed a gradual increase in burst duration with larger gCa, while a simultaneous decrease in burst frequency. Therefore, an increase in high-voltage activated Ca^2+^ current can account for the longer burst duration in noise-exposed LOC neurons.

We also tested the role of K2P (tonic) current and CDI in terminating bursts. **Figure 8B2** shows the spontaneous burst firing when tonic K^+^ conductance (gKLeak) was reduced to 0.03 mS/cm^2^, and there was a lengthening of burst duration. When gKLeak = 0, the model neuron fired tonically (data not shown). **Figure 8C2** shows the burst duration and frequency as a function of varying gKLeak. As gKLeak increased, burst duration decreased, highlighting the important role of tonic K^+^ current in truncating bursts. In contrast, the change in burst frequency was not monotonic: it initially increased and then decreased as K^+^ current increased. The increase in burst frequency was due to shorter burst duration, while the decrease in frequency was due to prolonged silent periods between bursts. When gKLeak exceeded 0.08 mS/cm^2^, the model neuron became completely silent.

To simulate the lack of CDI in the model neuron, we first calculated the weighted tau from tau 1 and tau 2 under four experimental conditions: EGTA and BAPTA, Ca^2+^ and Ba^2+^ (see Materials and Methods). This yielded the following four weighted tau values: Ca/EGTA (control) = 333 ms, Ca/BAPTA = 571 ms, Ba/EGTA = 954 ms, and Ba/BAPTA = 1328 ms. We then set the inactivation time constant of high-voltage activated Ca^2+^ current (tau_hLCa, also referred as *τ_h_* for *I*_Ca_ in Table 1) in the model to these values and measured burst duration and frequency. **Figure 8B3** shows the spontaneous burst firing when tau_hLCa was set to 1328 ms, simulating the elimination of CDI by both BAPTA and Ba^2+^. Burst duration gradually increased from 5.25 to 7.88 s with reduced CDI, suggesting CDI is important for burst termination (**Figure 8C3**). In a separate simulation, we set the Na^+^ current to 0 to examine how CDI regulates Ca^2+^-dependent oscillations in the model neuron (**Figure S3**). As expected, we observed a clear increase in burst duration as tau_hLCa increased, confirming that the facilitation of Ca^2+^ channel inactivation by CDI could contribute to termination of bursts. Based on these modeling data, we conclude that Ca^2+^ channel inactivation and tonic K^+^ current both contribute to burst termination in LOC neurons.

## Discussion

In this study, we demonstrated that Ca^2+^ channels are required for the spontaneous burst firing of LOC neurons and that Ca^2+^ current is modulated in an NIHL model. Ca^2+^ channels not only initiate bursts but also contribute to burst termination through their inactivation. NIHL significantly upregulates Ca^2+^ currents, resulting in elongated bursts, which suggests enhanced efferent activity within the noise-exposed cochlea. Additionally, NIHL reduces the role of CDI in facilitating inactivation, suggesting altered intracellular Ca^2+^ signaling in noise-exposed LOC neurons. These effects of NIHL on Ca^2+^ channels are specific to high-voltage activated channels, as T-type channels, though strongly expressed, were unaffected. Lastly, working synergistically with Ca^2+^ channel inactivation, K2P currents, including those mediated by TREK-1 channels, contribute to burst termination.

### Ionic mechanisms underlying spontaneous burst firing in LOC

Burst firing has been extensively studied in a wide variety of neurons. Besides bursts driven by alternating patterns of synaptic activity, the electrical interactions between ion channels within a single cell can serve as an intrinsic mechanism for spontaneous burst firing (35). A well-studied exemplar is the pancreatic β-cell; their alternating cycles of ATP/ADP ratios control the activation and deactivation of ATP-sensitive K^+^ channels, which play a fundamental role in bursts and the phasic release of insulin (36, 37). As in LOC neurons, the initiation of bursts in these secretory cells is driven by Ca^2+^ channels (36, 38). However, in other neurons, Na^+^ channels are instead crucial for burst initiation. For example, in pre-Botzinger complex neurons that control respiratory rhythm, persistent Na^+^ current is responsible for burst generation (39) In addition, we recently demonstrated that in fusiform cells – the principal cells of dorsal cochlear nucleus, persistent Na^+^ current and hyperpolarization-activated cyclic nucleotide-gated (HCN) channels work synergistically to generate membrane oscillations underlying spontaneous burst firing at a rate of ∼1 Hz (40).

In LOC neurons, Ca^2+^ channels, rather than persistent Na^+^ current, are crucial for burst generation, based on following evidence. In our previous study, TTX failed to abolish bursting activity assayed either with electrophysiology or calcium imaging (21). Furthermore, when Na^+^ channels and the majority of K^+^ channels were blocked, Ca^2+^ channels themselves were able to generate pace-making activity in current clamp simply by their alternating activation and inactivation cycles (see **Figure 5A**). Remarkably, the oscillation frequency of this pacemaker is not different from the burst frequency obtained without any channel antagonists (**Figure 5B**). In contrast to the predominant role of Ca^2+^ channels, Ca^2+^-activated or voltage-gated K^+^ channels appear less relevant in regulating bursts, as we saw no consistent effect of blockers of Ca^2+^-activated BK and SK, K_V_2, K_V_7, and other TEA-sensitive voltage-gated K^+^ channels on burst patterns. Since the K^+^ channels that contribute to burst termination are sensitive to Ba^2+^, we identified K2P and K_ir_ channels as potential candidates. K2P channels provide a tonic K^+^ leak conductance that could facilitate membrane repolarization when Ca^2+^ current is inactivated. And because LOC neurons have very high input resistances (∼1 GΩ) (21), a slight imbalance between outward and inward current could induce large swings in membrane voltage. This speculation is supported by our modeling data. When the K^+^ leak conductance was reduced, burst duration became longer, as a result of reduced repolarizing drive (see **Figure 8B2**). The role of K_ir_ channels is yet to be determined. Because K_ir_3.1 channels are strong inward rectifiers, we speculate that these channels might play a secondary role in burst termination (26). However, it is also possible that other types of K_ir_ channels are expressed at low levels in LOC neurons that are below detectability of snRNA-seq.

### Noise-induced hearing loss alters Ca^2+^ channel properties

LOC neurons express multiple types of Ca^2+^ channels that have differential susceptibility to NIHL: high-voltage activated Ca^2+^ current increases significantly one week following noise exposure while low-voltage activated current are little affected. The increased high-voltage activated Ca^2+^ current leads to prolonged bursting activity. It will be important to determine if this increase in current reflects an increased expression of ion channel density or a change in the kinetic properties of the channels that affects open probability. In addition, we found evidence suggesting a potential change in intracellular Ca^2+^ signaling. CDI is a unique feature of high-voltage activated Ca^2+^ channels, particularly L-type Ca^2+^ channels (30). In control neurons, manipulations such as using the fast Ca^2+^ chelator BAPTA or replacing external Ca^2+^ with Ba^2+^ attenuate CDI, significantly increased the time constants of inactivation. However, the effects of such manipulations are diminished in noise-exposed neurons, especially for the slow phase of inactivation (see **Figure 7B & 7C**). Among the reasons why CDI might be reduced in noise-exposed neurons is the possibility that expression of calmodulin could be downregulated by noise exposure. Calmodulin is an essential Ca^2+^-binding molecule required for CDI as well as for a variety of intracellular signaling cascades (30, 41, 42). Reduced CDI might also be due to an increased expression of Ca^2+^ binding proteins (CaBPs); indeed, expression of CaBP1, CaBP2 and CaBP4 is associated with the absence of CDI in mammalian inner hair cells, despite the expression of the L-type Ca^2+^ channel Ca_V_1.3 (43, 44). Reduced CDI could also result from post-transcriptional modifications. For example, Ca_V_1.3 in outer hair cells undergoes alternative splicing that results in a lack of calmodulin binding site (45).

It is noteworthy that noise-exposed LOC neurons still show prominent Ca^2+^ current inactivation even with compromised CDI properties, suggesting a parallel strengthening of voltage-dependent inactivation (VDI). The detailed mechanism of VDI in Ca^2+^ channels is still under debate but both α and auxiliary subunits play a role (30). In particular, β subunits are key modulators of VDI. Most of β subunit isoforms, except for β_2a_, promote VDI by either shifting VDI towards more hyperpolarizing voltages, or increasing inactivation kinetics (46). Mutants in β subunits serve as potential etiology for a variety of cardiovascular diseases (47). It will be interesting to examine whether the altered β subunits of Ca^2+^ channels are associated with NIHL.

### Downstream effects of enhanced LOC activity in the cochlea

LOC neurons express a variety of neurotransmitters, ranging from small molecules like acetylcholine, dopamine, and GABA, to larger peptide molecules including calcitonin gene-related peptide (CGRP), neuropeptide Y (NPY), urocortin, and possibly opioid peptides (14). Dopamine is inhibitory in the cochlea (20, 48), while CGRP is proposed to be excitatory (18). Urocortin might also regulate cochlear sensitivity and play a neuroprotective role for hearing during acoustic trauma (49, 50). Despite these previous studies, the precise effects and targets of synaptically-released neurotransmitters from LOC terminals on spiral ganglion neurons are unclear. What adds to this complexity is the biological consequence of increased bursting activity after noise exposure, as some of LOC’s neurotransmitters exert opposing effects (i.e., excitatory vs. inhibitory). Importantly, previous studies have demonstrated that the release of neuropeptides from synaptic terminals requires prolonged membrane depolarization provided by hundreds of presynaptic spikes (51, 52). Release was more effective if spike bursts were interleaved with silent intervals, which accommodates vesicle replenishment (53–56). Hence, longer bursts could be an optimized strategy to increase the amount of peptide release in the cochlea. In other words, NIHL might shift LOC’s preference from releasing small molecules under normal hearing conditions to a mixture of small molecules and neuropeptides. This is in line with higher peptide expression in noise-exposed LOC neurons (19). Increased CGRP in the cochlea could enhance the activity of spiral ganglion neurons, compensating for the reduced sound input, while increased urocortin might protect the auditory nerve fibers from acoustic trauma. Interestingly, many peptidergic neurons also exhibit seconds-long bursts similar to LOC neurons (37, 57, 58). In summary, the enhanced LOC activity in the cochlea might serve as an important compensatory strategy in response to NIHL, offering both compensation for reduced auditory input and protection for the cochlear function.

## Materials and Methods

### RESOURCE AVAILABILITY

#### Lead contact

Further information and resources should be directed to and will be fulfilled by the lead contact, Laurence Trussell.

#### Materials availability

This study did not generate new unique reagents.

#### Data and code availability

Electrophysiology and microscopy data reported in this paper will be shared by the lead contact upon request. MATLAB codes will be uploaded to GitHub upon publication.

### EXPERIMENTAL MODEL AND SUBJECT DETAILS

#### Mice

Mice were maintained in an animal facility managed by the Department of Comparative Medicine at Oregon Health & Science University. All procedures were approved by the Oregon Health & Science University’s Institutional Animal Care and Use Committee and met the recommendations of the Society of Neuroscience. Transgenic mice of both sexes expressing Cre recombinase under the endogenous choline acetyltransferase promoter (ChAT-IRES-Cre; Jackson Labs 031661) were used. These mice were crossed with a tdTomato reporter line (Ai9(RCL-tdT; Jackson Labs 007909) to generate mice that express tdTomato in cholinergic neurons (referred to as ChAT-Cre/tdTomato), including cholinergic lateral olivocochlear (LOC) neurons.

### METHOD DETAILS

#### Acute brainstem slice preparation

Juvenile and young adult mice at postnatal days (P) 17-34 were deeply anesthetized with isoflurane and decapitated. The brain was dissected in ice-cold (0°C) low-sodium artificial cerebral spinal fluid (aCSF) containing (in mM) 260 glucose, 25 NaHCO_3_, 2.5 KCl, 1.25 NaH_2_PO_4_, 1.2 CaCl_2_, 1 MgCl_2_, 0.4 ascorbic acid, 3 myo-inositol and 2 sodium pyruvate, continuously bubbled with a mixture of 95% O_2_/5% CO_2_. The brain was blocked coronally, affixed to the stage of a vibratome slicing chamber (VT1200S, Leica) and submerged in low-sodium aCSF. Coronal brain slices containing the superior olivary complex (SOC) were cut at 300 µm and transferred to normal aCSF at 34°C containing (in mM) 125 NaCl, 25 NaHCO_3_, 2.5 KCl, 1.25 NaH_2_PO_4_, 1.2 CaCl_2_, 1 MgCl_2_, 10 glucose, 0.4 ascorbic acid, 3 myo-inositol and 2 sodium pyruvate (pH 7.4, osmolarity 300-310 mOsm/l). Normal aCSF was continuously bubbled with a mixture of 95% O_2_/5% CO_2_. When sectioning was completed, slices were incubated for an additional 30 min at 34°C, followed by storage at room temperature, ∼22°C.

#### Electrophysiology

For electrophysiological experiments, slices were transferred to a recording chamber mounted on a Zeiss Axioskop 2 FS plus microscope. The microscope was equipped with a CCD camera, 10× and 40× water-immersion objectives. The recording chamber was perfused with normal aCSF at 3 ml/min and maintained at 31–33°C with an in-line heater (TC-324B; Warner Instrument Corp). Neurons in each slice were viewed using full-field fluorescence with a white-light LED attached to the epifluorescence port of the microscope that was passed through a tdTomato filter set. LOC neurons expressing tdTomato were identified in the lateral superior olive (LSO) that displayed a typical “S” shape on the coronal brainstem slice (21).

Borosilicate glass capillaries (OD 1.5 mm; World Precision Instruments) were pulled on a PP-830 Narishige vertical puller to a tip resistance of 3-6 MΩ for cell-attached and whole-cell voltage-clamp experiments, and of 6-10 MΩ for whole-cell current-clamp experiments. For cell-attached experiments, recording pipette was filled with normal aCSF. Whole-cell current-clamp experiments, except for the ones shown in **Figure 5A**, were performed with an internal pipette solution containing (in mM) 113 K-gluconate, 2.75 MgCl_2_, 1.75 MgSO_4_, 9 HEPES, 0.1 ethylene glycol tetraacetic acid (EGTA), 14 tris-phosphocreatine, 0.3 tris-GTP, 4 Na_2_-ATP, pH adjusted to 7.2 with KOH, and osmolality adjusted to 290 mOsm with sucrose. For whole-cell current-clamp experiments shown in **Figure 5A**, a CsCl-based internal solution was used, which contained (in mM) 103 CsCl, 10 tetraethylammonium chloride (TEA-Cl), 3.5 lidocaine N-ethyl chloride (QX-314-Cl), 2.75 MgCl_2_, 1.75 MgSO_4_, 9 HEPES, 0.1 EGTA, 14 tris-phosphocreatine, 0.3 tris-GTP, 4 Na_2_-ATP, with pH adjusted to 7.2 with CsOH, and osmolality adjusted to 290 mOsm with sucrose. To record K^+^ current, LOC neurons were patched with the aforementioned K^+^-based internal solution in the normal aCSF. To record Ca^2+^ current, whole-cell voltage-clamp experiments were conducted with a CsMeSO_3_-based internal solution containing (in mM) 103 CsMeSO_3_, 10 TEA-Cl, 2.75 MgCl_2_, 1.75 MgSO_4_, 9 HEPES, 0.1 EGTA, 14 tris-phosphocreatine, 0.3 tris-GTP, 4 Na_2_-ATP, with pH adjusted to 7.2 with CsOH, and osmolality adjusted to 290 mOsm with sucrose. LOC neurons were recorded in the external solution that contained (in mM) 100 TEA-Cl, 25 NaCl, 25 NaHCO_3_, 2.5 KCl, 1.25 NaH_2_PO_4_, 1.2 CaCl_2_, 1 MgCl_2_, 10 glucose, 0.4 ascorbic acid, 3 myo-inositol and 2 sodium pyruvate (pH 7.4, osmolarity 300-310 mOsm/l). In addition, 4-aminopyridine (4-AP, 3-4 mM) was included to block additional K^+^ current, especially A-type K^+^ current; NBQX (10 µM), MK-801 (10 µM), strychnine (0.5 µM) and gabazine (10 µM) to block excitatory and inhibitory synaptic activity; and TTX (0.5 µM) to block Na^+^ current. To attenuate Ca^2+^-dependent Ca^2+^ channel inactivation (CDI), either the extracellular 1.2 mM CaCl_2_ was replaced by 1.2 mM BaCl_2_, or a high-BAPTA internal solution was used, which contained (in mM) 63 CsMeSO_3_, 10 Cs_4_-BAPTA, 10 TEA-Cl, 2.75 MgCl_2_, 1.75 MgSO_4_, 9 HEPES, 14 tris-phosphocreatine, 0.3 tris-GTP, 4 Na_2_-ATP, with pH adjusted to 7.2 with CsOH, and osmolality adjusted to 290 mOsm with sucrose. In a subset of experiments (n = 6 cells), a NaHCO_3_- and NaH_2_PO_4_-free external solution was used to record Ca^2+^/Ba^2+^ current, containing (in mM) 100 TEA-Cl, 50 NaCl, 2.5 KCl, 10 HEPES, 1.2 CaCl_2_ or BaCl_2_, 1 MgCl_2_, 10 glucose, 0.4 ascorbic acid, 3 myo-inositol and 2 sodium pyruvate (pH 7.4 with NaOH, osmolarity 300-310 mOsm/l), along with the aforementioned drugs to block K^+^ and Na^+^ current, and synaptic receptors. This solution was continuously bubbled with 100% O_2_. Data acquired using two different external solutions were pooled and reported together. Recording pipettes were wrapped with Parafilm M (Bemis) to reduce pipette capacitance. Junction potential was experimentally measured as -10 mV, -3 mV and -10 mV for K^+^-, CsCl-, and CsMeSO_3_-based internal solution, respectively. All of our data were reported with the correction of junction potentials.

Electrophysiological experiments were performed using an Axon Multiclamp 700B amplifier (Molecular Devices). Raw data were low-pass filtered at 10 kHz and digitized at 100 kHz using a Digidata 1440A (Molecular Devices). Recording pipettes were visually guided to the LSO region and membrane patches were ruptured after a GΩ seal was attained. In cell-attached experiments, spontaneous burst firing activity was recorded for at least 5-10 minutes before applying various antagonists to the bath. In experiments using iberiotoxin (100 nM) or guangxitoxin (100 nM), bovine serum albumin (BSA, 0.5 mg/mL) was included in the bath, including the control aCSF, to prevent drug peptide absorption by the tubing. Iberiotoxin and guangxitoxin were allowed to circulate in the system for at least 10-15 minutes before subsequent recordings. In current-clamp experiments, LOC neurons were initially silent upon break-in, but all of them became spontaneously active within 5 minutes after whole-cell configuration. In some cases, up to -30 pA bias current was applied if a neuron became excessively depolarized. Passive membrane properties were measured by injecting a -5-pA current into the neuron for 1 s. Input resistance was calculated by dividing the injected current from the change in membrane voltage. The time constant (tau) was calculated by fitting a single exponential to the decay phase of the membrane voltage. Membrane capacitance was then calculated by dividing input resistance from tau. For all of the whole-cell voltage-clamp experiments, series resistance compensation was set to 70% correction and prediction. To record K^+^ current, LOC neurons were held at -70 mV. A series of 300-ms voltage steps were applied from -130 to +30 mV in a step of 10 mV. K^+^ currents in control conditions were monitored for at least 5 minutes before drug application to avoid any time-dependent rundown of the current. Leak subtraction was not applied to K^+^ current recordings. To record Ca^2+^ current, P/8 leak subtraction was applied as follows: individual LOC neurons were first held at -110 mV and then given a series of 500-ms voltage steps from -100 to +20 mV in a step of 10 mV. Second, holding voltage was changed to -70 mV and 2-s voltage steps from -70 to +10 mV were applied to the neuron. In some cases, we observed escaping Ca^2+^ current, as evidenced by a second, smaller peak in the recording, and those data were discarded. Inactivation kinetics were measured using Ca^2+^ current evoked by a single step from -70 to -20 mV for 2 s (holding voltage = -70 mV). A double exponential was fitted to the inactivation phase based on the following formula using the Simplex search method:

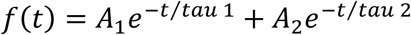

Tau 1 and tau 2 represents the decay time constant of fast and slow phase, respectively. Percent contribution of each tau to the total decay was calculated as:

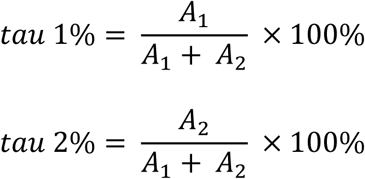

Then, weighted tau was calculated as:

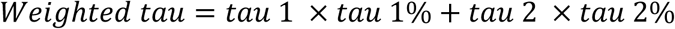

Finally, percent inactivation was calculated as percent reduction in Ca^2+^ current from the peak to the end of 2-s recording window.

#### Immunohistochemistry and confocal imaging

P25 mice were deeply anesthetized with isoflurane and then transcardially perfused with 0.1 M phosphate buffered saline (PBS) followed by 4% paraformaldehyde in 0.1 M PBS using a peristaltic pump. Brains were surgically extracted and incubated in 4% paraformaldehyde in 0.1 M PBS overnight at 4°C. On the next day, brains were rinsed three times in 0.1 M PBS, 10 minutes per rinse. Brains were then embedded in 4% agar and affixed to the stage of a vibratome slicing chamber and sectioned in the coronal plane at 50 µm. Each section was collected in 0.1 M PBS. For immunostaining, free-floating sections were permeabilized and blocked in 2% BSA, 2% fish gelatin, and 0.1% Triton X-100 in 0.1 M PBS for 2 hours at room temperature on a 2-D rocker. Sections were incubated in rabbit anti-TREK-1 (1:100, Alomone Labs) for 2 days at 4°C on a 2-D rocker, followed by three rinses in 0.1 M PBS, 10 minutes per rinse, then incubated in donkey anti-rabbit IgG Alexa Fluor 488 (1:500, Invitrogen) overnight at 4°C on a 2-D rocker. Sections were rinsed three times, 10 minutes per rinse on the next day, before being mounted on microscope slides and cover slipped with DAPI Fluoromount-G (Southern Biotech) mounting medium and sealed with clear nail polish. For negative control experiments, the primary antibodies were omitted, but all other steps remained the same. Low-magnification (20×) images of cerebellum were acquired on a Zeiss LSM900 confocal microscope, while high-magnification (63×) images of LOC neurons on a Zeiss LSM980 with Airyscan. All the images were processed for contrast and brightness in Fiji. LOC images in negative control experiments were captured using the same laser settings on the LSM980 and adjusted to similar levels of contrast and brightness in Fiji as the other LOC images.

#### Auditory brainstem response (ABR)

ChAT-Cre/tdTomato mice were anesthetized subcutaneously by a combination of 100 mg/kg ketamine and 10 mg/kg xylazine. Acoustic stimuli were digitally generated by a PXI data acquisition system with 24-bit analog-to-digital and digital-to-analog converter PXI-4461 card (National Instruments), amplified by SA1 stereo amplifier (Tucker-Davis Technologies), and delivered to the ear by a closed-field sound system consisting of two 15-mm speakers (CUI Devices CDMG15008-03A) and an electret microphone (Knowles FG-23329-P07) coupled to a probe tube. The probe tube was positioned over the ear canal and the speakers were calibrated using the probe tube microphone immediately prior to the experiment. ABRs were evoked by 5-ms tone pips at 4, 5.6, 8, 11.3, 16, 22.6 and 32 kHz. Sound levels were incremented in 5-dB steps from 10 to 90-dB SPL. The order of tone pip presentation was delivered in an interleaved manner that significantly reduced the ABR acquisition time (59). ABR signals were collected by three subcutaneous electrodes, inserted at vertex and along the ipsilateral mandible with ground near the base of the tail. 512 artifact-free averages were collected at each frequency and level to generate ABR waveforms for display. Hearing thresholds at each frequency were identified via visual inspection and defined as the lowest sound intensity at which wave 1 could be observed (60).

ChAT-Cre/tdTomato mice at P20-22 were tested with ABR, and only those with normal hearing were used in subsequent experiments. One day after the initial ABR tests, the mice were exposed to broadband noise at 110 dB SPL for 2 hours (see details below). 6 to 7 days after noise exposure, ABR tests were repeated to assess the degree of hearing loss. Brain slices were then collected within a week after the second ABR tests. Age-matched control mice were tested with ABR at P27-29 to ensure normal hearing. Brain slices from these mice were then used for experiments within a week.

#### Acoustic overexposure

Awake ChAT-Cre/tdTomato mice were placed unrestrained in small chambers within a subdivided cage (one mouse per chamber). The cage was positioned on a rotator in the center of a sound booth. Broadband noise (4-50 kHz) at 110 dB SPL was generated using custom software (61), amplified by a Crown D75A amplifier (Crown Audio), and delivered to the mice through an array of 9 horn tweeters (Fostex FT17H) mounted on 3 side walls of the sound booth (3 tweeters per side). A free-field ¼” microphone (PCB Piezotronics) was placed in the sound booth to monitor the intensity of the sound stimuli. The mice were exposed to noise for 2 hours.

#### Computational modeling of LOC neuron

A minimal, Hodgkin-Huxley-type model of LOC neuron was based on a previous model of peptidergic, burst firing subfornical organ neuron (62). This single-compartment model included a spherical soma with a 10-µm diameter and incorporated multiple ionic currents: *I*_Ca_, high-voltage activated Ca^2+^ current; *I*_CaT_, low-voltage activated Ca^2+^ current; *I*_Na_, transient Na^+^ current; *I*_NaP_, persistent Na^+^ current; *I*_K_, delayed-rectifier K^+^ current; *I*_A_, transient K^+^ current; *I*_KLeak_, tonic K^+^ current; *I*_KS_, slow-activating K^+^ current; *I*_NSCC_, non-selective cationic current (Ba^2+^-sensitive tonic inward current, see **Figure 3C1**, inset); *I*_L_, leak current, and *I*_noise_, an injected noise current. Input resistance is set at 1 GΩ. The membrane potential was described by:

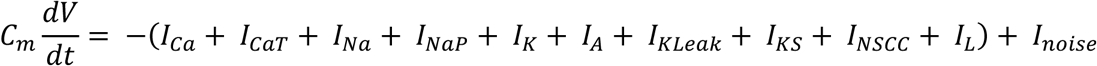

where *C*_m_ represents membrane capacitance in µF/cm^2^, calculated as the somatic capacitance divided by the soma area, with a 10-µm diameter. The somatic capacitance was measured as 5 pF for LOC neurons in whole-cell voltage-clamp mode. Consequently, *C*_m_ in the model was 1.59 µF/cm^2^ (62). *V* represents the membrane potential in mV, and *t* time in ms. *I*_noise_ represents Gaussian white noise with standard deviation of 2 µA/cm^2^. Other ionic currents are described by:

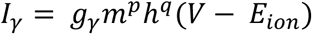

where *γ* refers to Ca, CaT, Na, NaP, K, A, KLeak, KS, NSCC or L. *g_γ_* represents maximal conductance of each ionic current in mS/cm^2^. *m* and *h* represent gating variables that determine the activation and inactivation of each channel, respectively. The corresponding *p* and *q* are the number of activation and inactivation gates for each channel. *E_ion_* is the reversal potential. *g*_L_ was calculated as 1/*R_m_* = 0.3183 mS/cm^2^, where *R_m_* = 3142 Ω * cm^2^. *g*_Ca_ and *g*_CaT_ were derived from our voltage-clamp recordings shown in **Figure 5D**. Other ionic conductances were adjusted accordingly to ensure the model neuron generated a spontaneous burst firing pattern with burst frequency and duration similar to the experimental data.

The gating variables *m* and *h* are further described by:

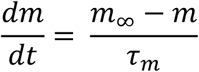

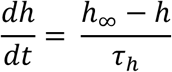

where *τ_m_* and *τ_h_* represent the activation and inactivation time constant of each ion channel, in ms. *τ_h_* for *I*_Ca_ (also referred as tau_hLCa) is set as weighted tau calculated above. In control, *τ_h_* = 333 ms, which was acquired with 0.1 mM EGTA in the pipette solution and 1.2 mM Ca^2+^ in the external solution. To simulate the attenuation of CDI, *τ_h_* was gradually increased to 571 ms (Ca/BAPTA), 954 ms (Ba/EGTA) and 1328 ms (Ba/BAPTA). *m_ꝏ_* and *h_ꝏ_* are steady-state activation and inactivation function, described by:

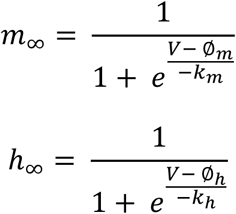

where *Ø_m_* and *k_m_* represent half activation voltage and the corresponding slope factor of *m*, while *Ø_h_* and *k_h_* half inactivation voltage and the corresponding slope factor of *h*. Simulations were performed in MATLAB (R2024a; MathWorks) and ordinary differential equations in this model were solved by Euler’s method with a time step of 0.01 ms. Numeric values of all model parameters are listed in Table 1.

#### Experimental design and statistical analysis

Electrophysiological data were collected using Clampex acquisition and Clampfit analysis software (version 10.9 and 11.3, Molecular Devices). GraphPad Prism 9 and 10 was used for statistical analysis and to make figures. Data are represented in the text and figures as mean ± SEM. Parametric analysis was used only after confirming assumptions of equal variances (F test for two-group comparison and Bartlett’s test for three-group comparison) and normality (D’Agostino-Pearson omnibus normality test). The equivalent nonparametric tests were used when data were not normal; the name of the test used is included with all statistics reported in the Results section. One-way ANOVA with repeated measures (RM) was used to compare statistics for single cells across more than two conditions, followed by Bonferroni *post hoc* comparisons. The Geisser and Greenhouse correction was applied to RM one-way ANOVAs to correct for possible violations of the assumption of sphericity (of note, this frequently changes associated degrees of freedom values to non-integers). Assumptions of sphericity and effective matching were also confirmed for one-way ANOVAs. Kolmogorov-Smirnov tests were performed to compare frequency distributions shown in **Figure 2A3-C3**. The standard for significant differences was defined as p < 0.05.

---

**Table S1.** KEY RESOURCES TABLE.

| **Reagent type (species) or resource** | **Designation** | **Source or reference** | **Identifier** |
| --- | --- | --- | --- |
| **Strain, strain background (M. musculus)** | ChAT-IRES-Cre | Jackson Laboratory<br>PMID: 21284986 | RRID: IMSR_JAX: 031661 |
| **Strain, strain background (M. musculus)** | Ai9(RCL-tdT) | Jackson Laboratory<br>PMID: 22446880 | RRID: IMSR_JAX: 007909 |
| **Strain, strain background (M. musculus)** | C57BL/6J | Jackson Laboratory | RRID: IMSR_JAX: 000664 |
| **Chemical compound, drug** | NBQX disodium salt | Tocris | Cat# 1044 |
| **Chemical compound, drug** | Tetrodotoxin (TTX) | Sigma Aldrich | Cat# 554412 |
| **Chemical compound, drug** | Nimodipine | Abcam | Cat# ab120138 |
| **Chemical compound, drug** | Strychnine hydrochloride | Sigma | Cat# S8753 |
| **Chemical compound, drug** | SR-95531 hydrobromide (gabazine) | Tocris | Cat# 1262 |
| **Chemical compound, drug** | (+)-MK-801 hydrogen maleate | Sigma Aldrich | Cat# M107 |
| **Chemical compound, drug** | Iberiotoxin | Alomone Labs | Cat# STI-400 |
| **Chemical compound, drug** | Apamin | Alomone Labs | Cat# STA-200 |
| **Chemical compound, drug** | Guangxitoxin-1E | Alomone Labs | Cat# STG-200 |
| **Chemical compound, drug** | XE-991 | Sigma Aldrich | Cat# X2254 |
| **Chemical compound, drug** | Tetraethylammonium chloride (TEA-Cl) | Sigma Aldrich | Cat# 86614 |
| **Chemical compound, drug** | 4-aminopyridine (4-AP) | Sigma Aldrich | Cat# 275875 |
| **Antibody** | Anti-TREK-1 (rabbit polyclonal) | Alomone Labs | Cat# APC-047<br>RRID: AB_2040136 |
| **Antibody** | Anti-rabbit Alexa Fluor 488 (donkey polyclonal) | Invitrogen | Cat# A-21206<br>RRID: AB_2535792 |
| **Software, algorithm** | pClamp 10 and 11 | Molecular Devices | RRID: SCR_011323 |
| **Software, algorithm** | MATLAB R2024a | MathWorks | RRID: SCR_001622 |
| **Software, algorithm** | ZEN Blue | Zeiss | RRID: SCR_013672 |
| **Software, algorithm** | Fiji | Fiji.sc | RRID: SCR_002285 |
| **Software, algorithm** | Excel | Microsoft | RRID: SCR_016137 |
| **Software, algorithm** | GraphPad Prism 9 and 10 | GraphPad | RRID: SCR_002798 |
| **Software, algorithm** | Affinity Designer | Serif | RRID: SCR_016952 |
| **Software, algorithm** | Auditory Wave Analysis (0.0.2) | Zenodo | (60) |
| **Software,**<br>**algorithm** | psiexperiment/noise-<br>exp: v0.0.1 (v0.0.1) | Zenodo | (61) |

## Acknowledgments

This work was supported by NIH grants DC004450 and NS116798 to L.O.T. We thank members of the Trussell lab and Dr. Brad Buran for helpful discussions.

## Notes

### Competing Interest Statement

The authors have declared no competing interest.

