## Supplemental Figures 1-3 for "Noise-induced hearing loss enhances Ca^2+^-dependent spontaneous bursting activity in lateral cochlear efferents"

Supplemental Figures 1-3 for Hong and Trussell “**Noise-induced hearing loss enhances Ca2+-dependent spontaneous bursting activity in lateral cochlear efferents**”

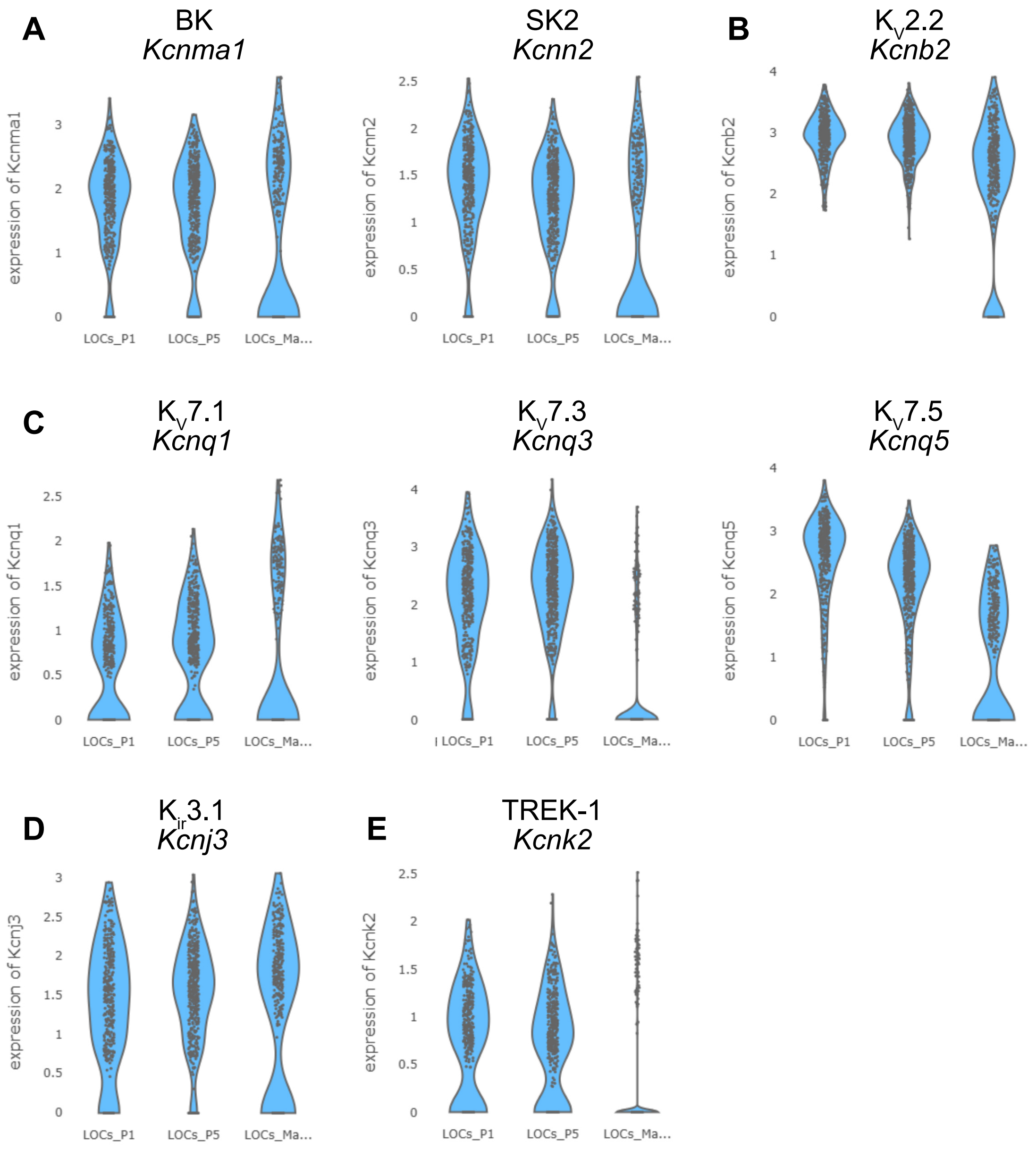
A
BK
Kcnma1
SK2
Kcnn2
B
KV2.2
Kcnb2
C
KV7.1
Kcnq1
KV7.3
Kcnq3
KV7.5
Kcnq5
D
Kir3.1
Kcnj3
E
TREK-1
Kcnk2
expression of Kcnma1
expression of Kcnn2
expression of Kcnb2
expression of Kcnq1
expression of Kcnq3
expression of Kcnq5
expression of Kcnj3
expression of Kcnk2
LOCs_P1
LOCs_P5
LOCs_Ma...

Fig S1

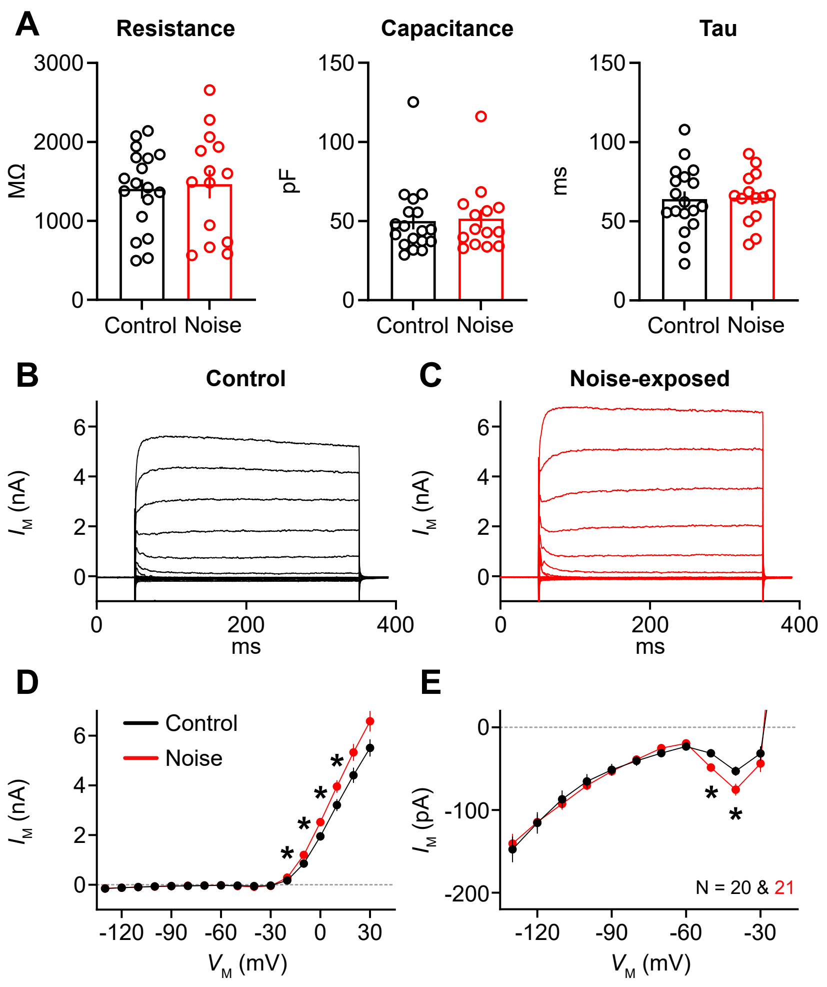
A
Resistance
Capacitance
Tau
MΩ
pF
ms
Control Noise
B
Control
C
Noise-exposed
$I_M$ (nA)
ms
D
Control
Noise
$V_M$ (mV)
E
$I_M$ (pA)
N = 20 & 21

Fig S2

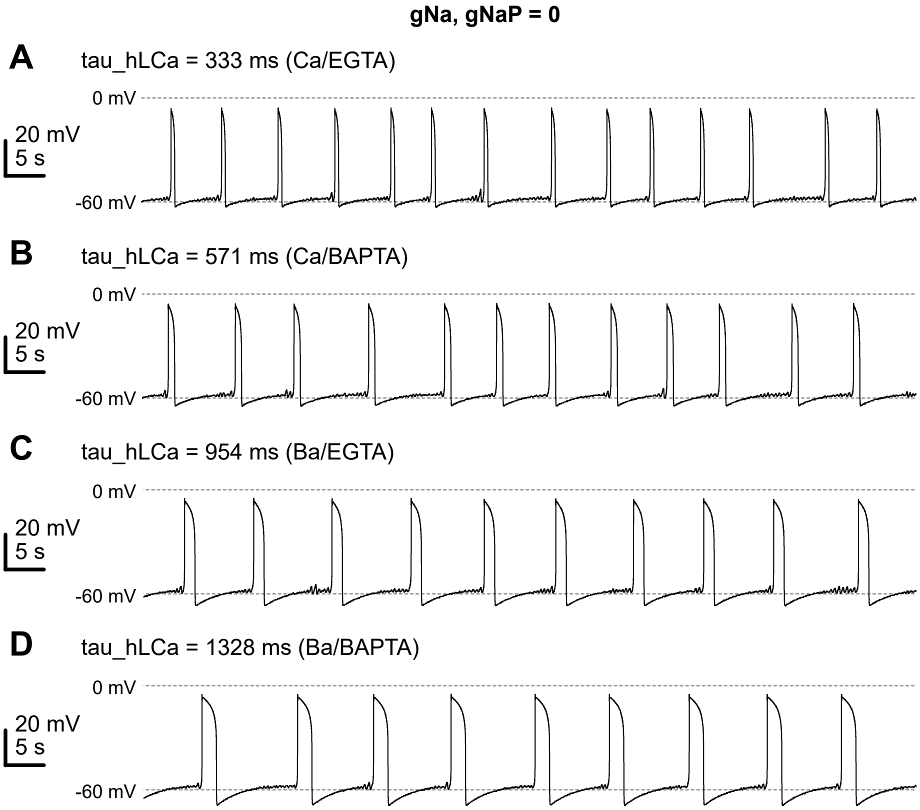
gNa, gNaP = 0
A
tau_hLCa = 333 ms (Ca/EGTA)
0 mV
20 mV
5 s
-60 mV
B
tau_hLCa = 571 ms (Ca/BAPTA)
0 mV
20 mV
5 s
-60 mV
C
tau_hLCa = 954 ms (Ba/EGTA)
0 mV
20 mV
5 s
-60 mV
D
tau_hLCa = 1328 ms (Ba/BAPTA)
0 mV
20 mV
5 s
-60 mV

Fig S3
